# Specification of Origins of DNA Replication Within Distinct Chromatin States in Human Cells

**DOI:** 10.64898/2026.10.22.761122

**Authors:** Meng Ouyang, Bruce Stillman

**Affiliations:** Cold Spring Harbor Laboratory, 1 Bungtown Road, Cold Spring Harbor, NY 11724, USA

**Keywords:** Origins of DNA replication, Replication timing, Chromatin 3D structures, Chromatin States, CTCF

## Abstract

A comprehensive map of binding sites for the Origin Recognition Complex (ORC) in the human genome reveals relationships among replication licensing, chromatin state, three-dimensional genome organization, location of origins of DNA replication and DNA replication timing. Defining these relationships is essential for understanding how replication licensing is integrated with the broader organization and duplication of the human genome. Using 30 high-quality ORC1 CUT&RUN datasets, an ORC1 three-dimensional binding matrix (ORC1 3DM) was constructed that captures genomic position, binding intensity, and reproducibility across replicates. This analysis reveals considerable heterogeneity in ORC1 binding, and integration with chromatin-state maps shows that ORC1-binding sites separate into distinct epigenomic groups, with high-confidence sites enriched in H2A.Z1- and H3K4me3-associated open chromatin. Hi-C analysis demonstrates preferential localization of ORC1 binding to sub-regions of genomic A compartments and frequent association with loop anchors, supported by a direct interaction between ORC1 and CTCF. Replication-timing analysis shows that ORC1-binding sites are enriched in initiation zones and exhibit subgroup-specific temporal patterns. An integrated analysis combining multiple replication initiation assays and chromatin features further places ORC1 3DM within a unified replication–chromatin framework. Together, these results support a hierarchical, probabilistic model of replication licensing that is embedded within chromatin architecture.

**Significance Statement:** Understanding how DNA replication licensing relates to chromatin organization requires quantitative mapping and integration of genomic features. We developed an ORC1 three-dimensional binding matrix (ORC1 3DM) from 30 CUT&RUN datasets, quantifying genomic binding coordinates, intensity and reproducibility. We established an integrative framework to compare diverse genomic datasets across 25 chromatin states and a high-resolution approach to characterize replication timing profiles and their relationships with replication initiation. These approaches revealed distinct chromatin-associated ORC1-binding groups, preferential localization of high-confidence ORC1 sites in active chromatin and highly interactive A-compartment subregions, and subgroup-specific relationships with replication initiation and timing. Together, these findings and methodological advances provide a quantitative framework for investigating relationships among replication licensing, chromatin state, three-dimensional genome organization, and replication timing.

## Introduction

Eukaryotic genomes are large, multi-chromosomal polymers of DNA and protein that must be duplicated precisely once per cell cycle. Because the length of individual chromosomes far exceeds the distance that a single pair of replication forks can traverse within S phase, genome duplication relies on the activation of many replication origins that are distributed along each chromosome. How replication origins are defined, positioned, and regulated to ensure complete and timely genome duplication remains a fundamental question in eukaryotic cell biology, particularly in the human genome (1–4).

In the budding yeast *Saccharomyces cerevisiae*, replication origins have been characterized in the greatest detail (5, 6). Origins are specified by discrete DNA elements that are recognized by the Origin Recognition Complex (ORC) in a DNA sequence-dependent manner, as well as the intrinsic ability of the origin DNA to bend upon binding ORC (7–15). ORC binding during mitotic exit and G1 phase initiates the ordered assembly of pre-Replicative Complexes (pre-RCs), which includes the recruitment of Cdc6, followed by the Cdt1-mediated loading of the Mcm2–7 helicase precursor as a head-to-head double hexamer (16). This licensing step establishes a pool of potential origins but only a subset is activated throughout S phase to generate bidirectional replication forks and complete replication of each chromosome (6, 14, 17–20).

Outside the budding yeast *S. cerevisiae*, origin specification appears more diverse (1, 2, 14, 21). The budding yeast *Yarrowia lipolytica* has a very loose DNA consensus sequence that contributes to origin specification and unlike *S. cerevisiae*, requires both ORC and Cdc6 for origin recognition (22, 23). In fission yeast *Schizosaccharomyces pombe* and several other fungal species, origin activity is associated with broader, AT-rich regions rather than discrete consensus sequences (24). Studies in organisms such as *Drosophila* and *Caenorhabditis elegans* further indicate that chromatin context, transcriptional activity, and higher-order genome organization contribute to origin selection (25–35). These observations suggest that while the core replication machinery is conserved, the mechanisms defining origin location have diverged substantially across eukaryotes.

In human cells, multiple assays have been developed for locating origins of DNA replication, including Bubble-seq (36), Ini-seq (37, 38), OK-seq (4, 39), SNS-seq (40) and optical replication mapping (41) for initiation-site detection, but a consensus, high resolution map of human origins remains elusive. Furthermore, the determinants of replication origin recognition and specification remain incompletely defined. Biochemical studies indicate that human ORC binds DNA with limited DNA sequence specificity (42–45), and genome-wide analyses have failed to identify a universal origin consensus sequence. While pre-RC assembly by ORC, CDC6, CDT1 and MCM2-7 on DNA has been reconstituted with purified human proteins (46–48), these studies have not addressed how origins are located in vivo. Nevertheless, human cells exhibit highly reproducible replication timing programs, implying that origin usage is constrained by stable genomic features rather than being purely stochastic (2, 49–55). A number of efforts have been made to map ORC-binding sites in the human genome using approaches such as sheared chromatin immunoprecipitation and DNA sequencing (ChIP-seq) (56–59), but these studies show very limited concordance between studies and in some cases, even low overlap between different ORC protein subunits, reflecting either technical challenges, biological complexity, or both. More importantly, the identified binding sites show limited overlap with known sites of initiation of DNA replication, which is entirely unexpected (56, 60, 61). Thus, it was not clear whether ORC specifies the locations of origins of DNA replication in the human genome.

In this study, we sought to generate a comprehensive, quantitative map of ORC1 binding sites in the human genome. To achieve this, we employed CUT&RUN, a sensitive method for mapping protein–DNA interactions (62, 63), and integrated ORC1 profiling data across multiple replicates to construct an ORC1 three-dimensional binding matrix (ORC1 3DM) that captures genomic location, binding intensity, and reproducibility. To investigate ORC1 binding within a broader genomic context, we integrated the ORC1 3DM with diverse genomic datasets. These analyses revealed relationships between ORC1 binding and the binding of other ORC subunits, pre-RC proteins such as CDC6, MCM2 and MCM3, and ORC-interacting proteins, as well as association with known DNA replication initiation sites, chromatin states, three-dimensional genome organization, and DNA replication timing patterns. Together, our results provide an integrated view of the genomic distribution of replication licensing, its relationships with higher-order chromatin and chromosome architecture, and its strong association with DNA replication initiation landscapes.

## Results

### High-quality ORC1 CUT&RUN data reveal reproducible and heterogeneous binding patterns

CUT&RUN was used to map ORC1 binding genome wide. The resulting datasets displayed sharp, reproducible peaks using an anti-ORC1 antibody, with minimal background observed using an IgG control (**Fig. 1A and 1B**). Peaks were called in two modes, lambda and non-lambda mode using macs2 software (64). This analysis resulted in 36,016, 34,538 and 83,801 peaks from ORC1 replicates 1-3 compared to 49 peaks from the IgG control in lambda mode; and in the non-lambda mode, 111,823, 159,115 and 280,293 peaks from ORC1 replicates 1-3 and 492 peaks from the IgG control. The variation in peak numbers among replicates reflects differences in sequencing depth as well as the distinct background estimation strategies implemented in MACS2, namely the local background model (lambda mode) and the global background model (non-lambda mode). This suggests that it is not possible to rely solely on peak calling to determine the data quality for this type of experiment. Nevertheless, the significant differences in peak numbers between ORC1 and IgG are obvious. High-confidence peaks were identified in each replicate, with Fraction of Reads in Peaks (FRiP) scores of 13.1%∼25% in ORC1 libraries, while 0.1% in the IgG control library, indicating strong enrichment. Correlation analysis shows a good correlation between ORC1 replicates and low or no correlation between IgG and ORC1 (**Fig. 1C and S1A**).

**Fig. 1.**
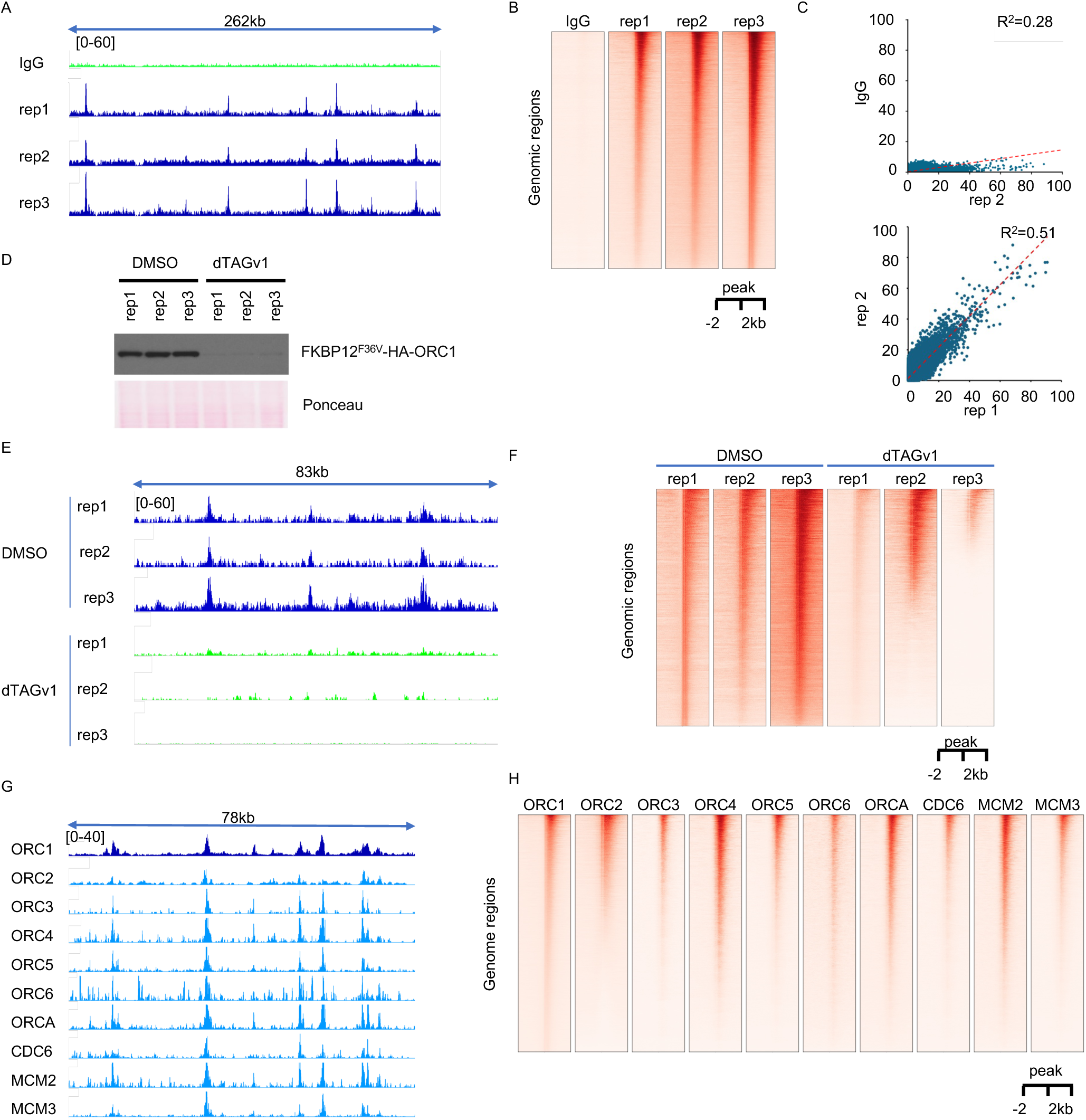
Genome-wide mapping and specificity of ORC1 binding. (A) ORC1 (Bethyl, A301-892A) and IgG CUT&RUN tracks in U2OS cells at chr2:17,960,000–18,222,000. (B) Corresponding signals within 2 kb of ORC1 peak summits. (C) Read-count comparisons on chromosome 21 between IgG and ORC1 replicate 2 and between ORC1 replicates 1 and 2; displayed values are R². (D) Immunoblot of endogenous FKBP12F36V-tagged ORC1 after DMSO or 100 nM dTAGv1 for 2 h; Ponceau staining shows loading. (E and F) Spike-in-normalized ORC1 CUT&RUN signals after the same treatments, shown at chr2:66,822,819–66,906,107 (E) and around reference ORC1 peak summits (F). Reference peaks were defined in DMSO replicate 1. (G and H) Tracks at chr10:33,206,637–33,285,430 (G) and heatmaps centered on ORC1 peak summits (H) for ORC1, ORC2, ORC3–ORC6, ORCA, CDC6, MCM2, and MCM3. ORC1 and ORC2 were mapped by CUT&RUN; the other proteins were mapped by CUT&Tag. Replicate labels identify biological replicates. Bracketed values indicate track display ranges.

A general concern for sequencing-based chromatin profiling methods such as ChIP-seq, CUT&RUN, and CUT&Tag is that apparent enrichment can be confounded by genomic mapping biases, such as enrichment for genome accessible sites (65, 66). To test whether the ORC1 peaks we observed represent genuine binding rather than artifacts, we employed an ORC1-degron approach. Using CRISPR, the endogenous ORC1 was tagged at its N-terminus with an FKBP12^F36V^ degron to generate a stable cell line (**Fig. S1B**) (67). We obtained three cell-line clones. Time course experiments show that the tagged endogenous ORC1 proteins were very sensitive to dTAGv1 treatment, observing a significant decrease in ORC1 levels after treatment with 100nM dTAGv1 for 1hr, and a further decrease with prolonged treatment (**Fig. S1C**).

Although prolonged treatment can achieve higher depletion, it may introduce secondary phenotypes or effects. As a trade-off, a depletion for 2 hr was selected for subsequent experiments. At this time point, ORC1 levels were nearly depleted (**Fig. 1D**), while cells showed no observable phenotypic abnormalities by microscopy. CUT&RUN analysis of DMSO-(control) and dTAGv1-treated cells revealed that both peak intensity and global signal distribution were dramatically reduced upon ORC1 depletion, confirming that the CUT&RUN signal was ORC1-dependent and highly specific (**Fig. 1E and 1F**).

Based on studies in *S. cerevisiae* and biochemical studies in vitro with purified human proteins, replication licensing requires the ordered assembly of multiple proteins to form pre-Replicative Complexes (pre-RC) that mark sites of initiation of DNA replication (14–16). To assess co-localization among components of the ORC, we profiled additional ORC subunits (ORC2–6) together with CDC6, ORCA (an ORC associated protein, also called LRWD1 (68)) and MCM2 and MCM3 using CUT&RUN or CUT&Tag. The Integrative Genomics Viewer (IGV) (69) tracks show that all these proteins follow a similar binding pattern as ORC1 (**Fig. 1G**). Heatmaps centered on ORC1 peaks reveal that ORC4, ORCA, and MCM2 exhibit the strongest co-localization with ORC1, whereas ORC2 shows the weakest overlap and the remaining subunits display intermediate levels (**Fig. 1H**). This hierarchy most likely reflects the use of different antibodies in the experiments and their ability to access the chromatin-bound proteins. Thus, inherent limitations of the CUT&RUN and CUT&Tag assays, such as differences in antibody efficiency, should be considered when interpreting these patterns. It is also possible that different ORC subunits and pre-RC proteins have individual functions in addition to pre-RC assembly since it is known that ORC has functions in addition to those in DNA replication (30, 70–73). Nevertheless, the co-localization of all ORC subunits with CDC6 and MCM proteins is consistent with the idea that, as in yeasts, these proteins cooperate to specify sites of initiation of DNA replication.

## Construction of an ORC1 three-dimensional binding matrix (ORC1 3DM)

Despite the apparent reproducibility observed in IGV tracks, a quantitative comparison revealed variable overlap in binding sites between ORC1 CUT&RUN replicates in a single experiment (**Fig. 2A**). This discrepancy arises partly from limitations of peak-calling algorithms, which are sensitive to signal intensity and local background variations, but also reflects biological and technical variability inherent to single-replicate assays. Similar low-overlap phenomena have been reported in other CUT&RUN studies (74), suggesting that such variation is a general property of these techniques rather than an artifact of individual experiments. To minimize this variation, we adopted two complementary strategies. First, we sought to standardize our CUT&RUN protocol as thoroughly as possible. Although this was challenging due to the many experimental steps involved, careful optimization of each stage helped reduce technical variability. Second, we increased the number of biological replicates by generating many independent libraries. In total, we generated 44 ORC1 CUT&RUN libraries and based on two quality metrics, i.e. FRiP score and minimum peak number, 30 high-quality libraries were selected for downstream analysis. To systematically integrate these data, we developed a computational framework (SI Appendix, Material and Methods, **Fig. S2A**) to construct a three-dimensional ORC1-binding site matrix (ORC1 3DM), which characterizes each genomic bin by its coordinate, binding intensity, and reproducibility (or confidence measured as frequency of detection across replicates).

**Fig. 2.**
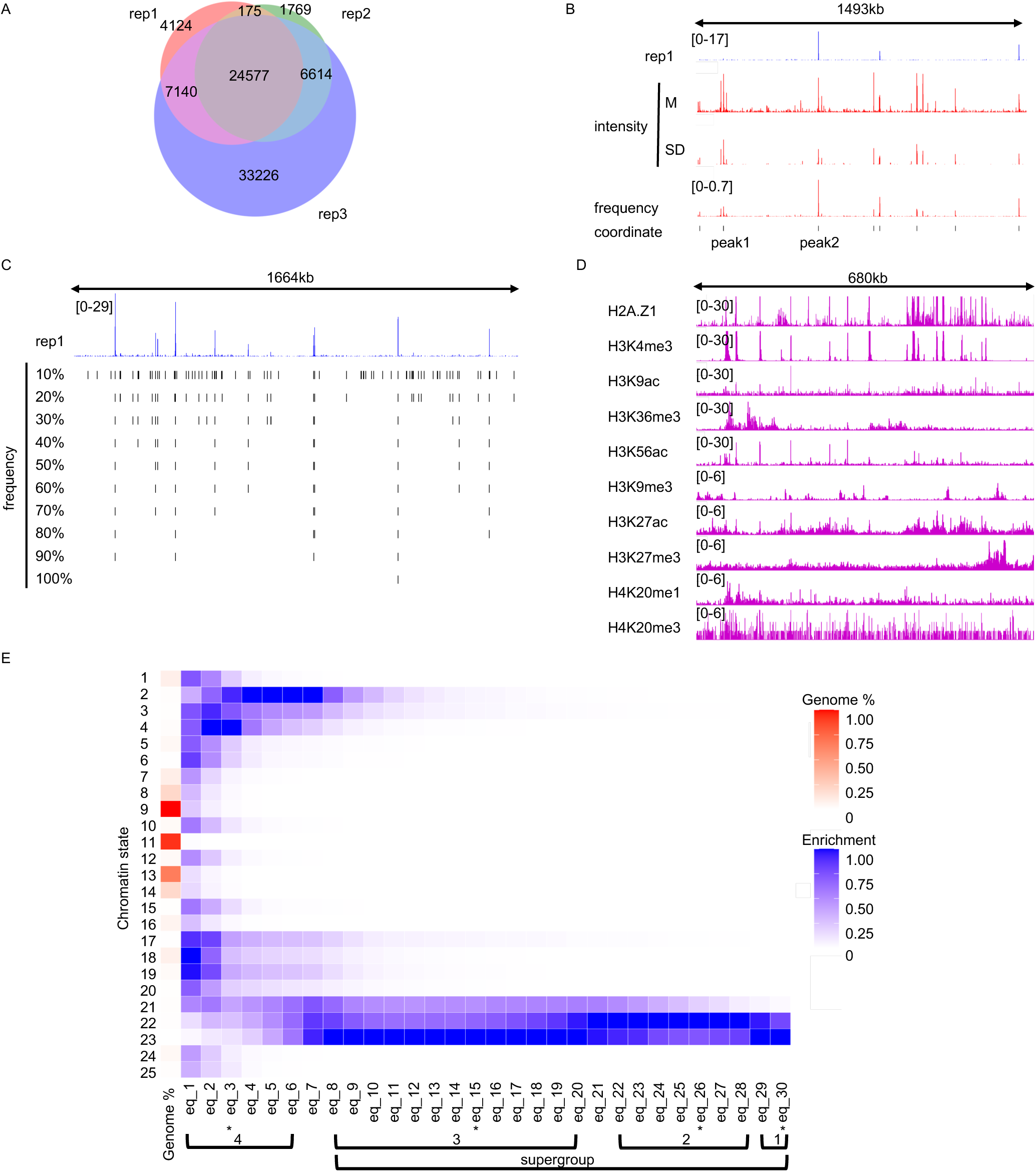
Construction and chromatin-state stratification of the ORC1 three-dimensional binding matrix (ORC1 3DM). (A) Overlap of ORC1 CUT&RUN peaks from three biological replicates. (B) Representative ORC1 3DM tracks at chr2:137,952,630–139,446,079. M: mean; SD: standard deviation. Peaks 1 and 2 illustrate distinct intensity and frequency patterns. (C) Frequency-threshold views of the ORC1 3DM at chr2:38,222,164–39,886,372. Frequency is the fraction of the 30 biological replicates in which a site was detected. (D) Tracks for ten histone profiles at chr5:139,212,636– 139,893,404. H2A.Z1, H3K4me3, H3K9ac, H3K9me3, H3K27ac, and H3K27me3 were mapped by CUT&RUN; H3K36me3, H3K56ac, H4K20me1, and H4K20me3 were mapped by CUT&Tag. (E) Relative genomic coverage and ORC1 enrichment across the 25 chromatin states. eq_n denotes sites detected in exactly n of 30 biological replicates. Genomic coverage is normalized by its maximum across states, and ORC1 enrichment is independently normalized by its maximum within each frequency-subset column. Asterisks identify eq_30, eq_26, eq_15, and eq_3, representing groups 1–4, respectively, in Fig. 5.

To visualize the integrated data structure, the ORC1 3DM on IGV was displayed as a multi-track summary view (**Fig. 2B**). From top to bottom, the tracks represent: (1) an ORC1 CUT&RUN replicate, which served as the reference; (2) the non-zero averaged binding intensity across the 30 selected libraries; (3) the non-zero standard deviation of the binding intensity; (4) frequency; and (5) the genomic coordinate. This visualization highlights both the reproducibility and the variability of ORC1 binding across experiments. Two representative examples, *peak 1* and *peak 2*, illustrate this approach: although their averaged signal intensities are comparable, their detection frequencies differ substantially, with *peak 1* being low-frequency and *peak 2* high-frequency. ORC1 3DM therefore provides a more comprehensive description of each binding event. It enables selective filtering of peaks based on the multiple parameters, e.g. frequency and intensity, which is valuable for downstream analyses. To demonstrate one practical use of ORC1 3DM, we stratified peaks by every 10% detection frequency (**Fig. 2C**). This frequency-based slicing approach allows the systematic comparison of subsets with varying confidence levels, facilitating the identification of high-confidence, consistently bound sites as well as variable, context-dependent regions. The distribution of intensity of the peaks that have different frequencies across each experiment is shown in **Fig. S2B,** demonstrating that more conserved peaks tend to have higher signal intensity. Given that the total collection of ORC1 peaks is not homogeneous but instead exhibits a hierarchical organization, obtaining an accurate representation of the distribution of ORC1 binding sites in the human genome requires many replicates and an analysis such as ORC1 3DM.

## Chromatin-state stratification of ORC1 3DM reveals heterogeneous groups of ORC1-binding sites

To further dissect the heterogeneity of ORC1 peaks, we stratified ORC1 3DM by chromatin states. To this end, we generated CUT&RUN or CUT&Tag libraries for ten different histone marks representing key regulatory and structural chromatin features: H2A.Z1, H3K27ac, H3K27me3, H3K36me3, H3K4me3, H3K56ac, H3K9ac, H3K9me3, H4K20me1, and

H4K20me3. Together, these marks span promoter, enhancer, gene-body, transcribed, open, and closed chromatin regions (75). For each histone modification, at least three biological replicates were produced, except H3K56ac, for which two were produced, with two or more showing high reproducibility. For simplicity, we selected one replicate with the highest FRiP score per histone mark for downstream analysis (**Fig. 2D and S2C**). ChromHMM (76, 77), a widely applied chromatin-segmentation algorithm, was used to train multistate models using combinations of these histone marks. Models with 10, 15, 20, 25, and 30 states were tested, and the 25-state model was chosen as the optimal balance between resolution and interpretability. The resulting states are well separated and correspond to distinct functional elements such as active promoters, enhancers, transcribed regions, and repressive domains (**Fig. S2E and S2F**). We then divided the ORC1 3DM into 30 subsets (eq_1∼30) according to their reproducibility. For example, eq_30 includes peaks present in all 30 libraries, and eq_10 includes peaks present in 10 libraries. Applying the 25-state chromatin model to these subsets revealed a clear hierarchical organization of ORC1 binding (**Fig. 2E**). From high to low frequency along the x-axis, the subsets aggregated into four major groups. Group 1 (eq_29∼30) was characterized by strong enrichment in chromatin state 23, followed by state 22; Group 2 (eq_22∼28) was enriched primarily in state 22, with secondary enrichment in states 21 and 23; Group 3 (eq_8∼20) was similar to Group 1 but with additional enrichment for state 21 and finally, Group 4 (eq_1∼6) was mainly enriched in states 2∼4 with broader coverage across states 1∼6 and 17∼23. Intermediate subsets (eq_7 and eq_21) exhibited transitional patterns between neighboring groups. Groups 1∼3 were all dominated by states 21∼23 and were distinct from Group 4 and therefore could be collectively described as a supergroup.

Of note, when we examined the histone mark compositions of the supergroup (Groups 1∼3) and Group 4, the most striking difference was H2A.Z1, which was highly enriched in states 21∼23 but nearly absent in states 2∼6 (**Fig. S2D**). This contrast suggests that H2A.Z1 may facilitate ORC1 association with DNA and increase its detectability in CUT&RUN assays. All four groups showed high levels of H3K4me3, indicating that ORC1 binding generally occurred in promoter-proximal or transcriptionally permissive regions. The supergroup (states 21∼23) shared a core composition of H3K4me3 and H2A.Z1, while differences among these states were defined by two additional histone-mark clusters (**Fig. S2D**): Cluster 1: H3K36me3, H4K20me1, H4K20me3;

Cluster 2: H3K56ac, H3K27ac, H3K9ac. State 23 showed high levels of both clusters, corresponding to the most transcriptionally active chromatin. State 22 was enriched for Cluster 2 but low in Cluster 1, consistent with open, acetylated regions lacking elongation-associated marks. State 21 showed low levels of both clusters, suggesting a more poised or less active chromatin context. These chromatin features revealed a clear hierarchy among ORC1-binding groups. The supergroup (Groups 1∼3) lay in open, transcriptionally active regions enriched for H2A.Z1 and H3K4me3, environments generally associated with high nucleosome turnover and accessibility. In contrast, Group 4, linked to states 2∼4 and their neighbors, occupied more compact chromatin with low but detectable H3K9me3 and H3K27me3, distinct from the strongly repressive states (12∼17, 20). Notably, states 21∼23 lacked H3K9me3 and showed only low-to-moderate H3K27me3, indicating a largely non-repressive environment. Collectively, the ORC1 binding sites represent a small proportion of the human genome (**Fig. 2E**). Together, these results highlight that ORC1 binding occurred across a broad chromatin spectrum—from highly active, accessible regions to weakly repressive domains - reflecting a structured hierarchy of chromatin states with potential functional diversity in DNA replication licensing/initiation or non-replication functions.

## ORC1 3DM correlates with higher-order chromatin 3D structure

Chromatin three-dimensional (3D) architecture is a major regulator of nuclear processes, including DNA replication (78). To investigate how ORC1 binding relates to genome organization, we generated Hi-C data (79) from the same cell line. The resulting contact matrices revealed characteristic multiscale structural features, including A and B compartments that define loosely and tightly packed chromatin, respectively, topologically associating domains (TADs), and chromatin loops (**Fig. 3A**). ORC1 3DM peaks preferentially localized to A compartments, with 62.4% in A compartments and 37.6% in B compartments (**Fig. 3A and 3B**), consistent with their association with active and accessible chromatin. This preference was more pronounced for the ORC1 3DM supergroup, with coverage-normalized proportions of 83.6% in A compartments and 16.4% in B compartments (**Fig. S3A**). Examination of the Hi-C eigenvector revealed substantial variation in compartment signal within both A and B compartments. To characterize this heterogeneity, we used Gaussian mixture modeling of compartment eigenvector values to subdivide the A and B compartments into four sub-compartments each (A1–A4 and B1–B4). These sub-compartments represent gradients of Hi-C compartment signal, with A1 showing the strongest positive signal and A4 the weakest, and B1– B4 forming a corresponding gradient within B compartments (**Fig. S3B**). Mapping ORC1 3DM peaks onto these sub-compartments revealed that A2 and A3 together contained the majority of ORC1-binding sites in the supergroup **(Fig. 3C and S3E**). However, the sub-compartments differed substantially in genomic coverage (**Fig. S3D**). After normalizing for genomic coverage, ORC1 peaks were enriched in all A sub-compartments, with fold enrichment decreasing in the order A1 > A2 > A3 > A4, whereas all B sub-compartments showed depletion, most prominently in B2 and B3 and less strongly in B4 (**Fig. S3E**). Further analysis restricted to A compartments, with positive log₂ enrichment scores normalized within each peak group, identified A1 as the largest contributor to positive enrichment, followed by A2 (**Fig. S3F**). Together, these results demonstrate that ORC1 binding is nonuniformly distributed within A/B compartments: although A2 and A3 contain most ORC1-binding sites in the supergroup, A1 exhibits the strongest enrichment relative to genomic coverage.

**Fig. 3.**
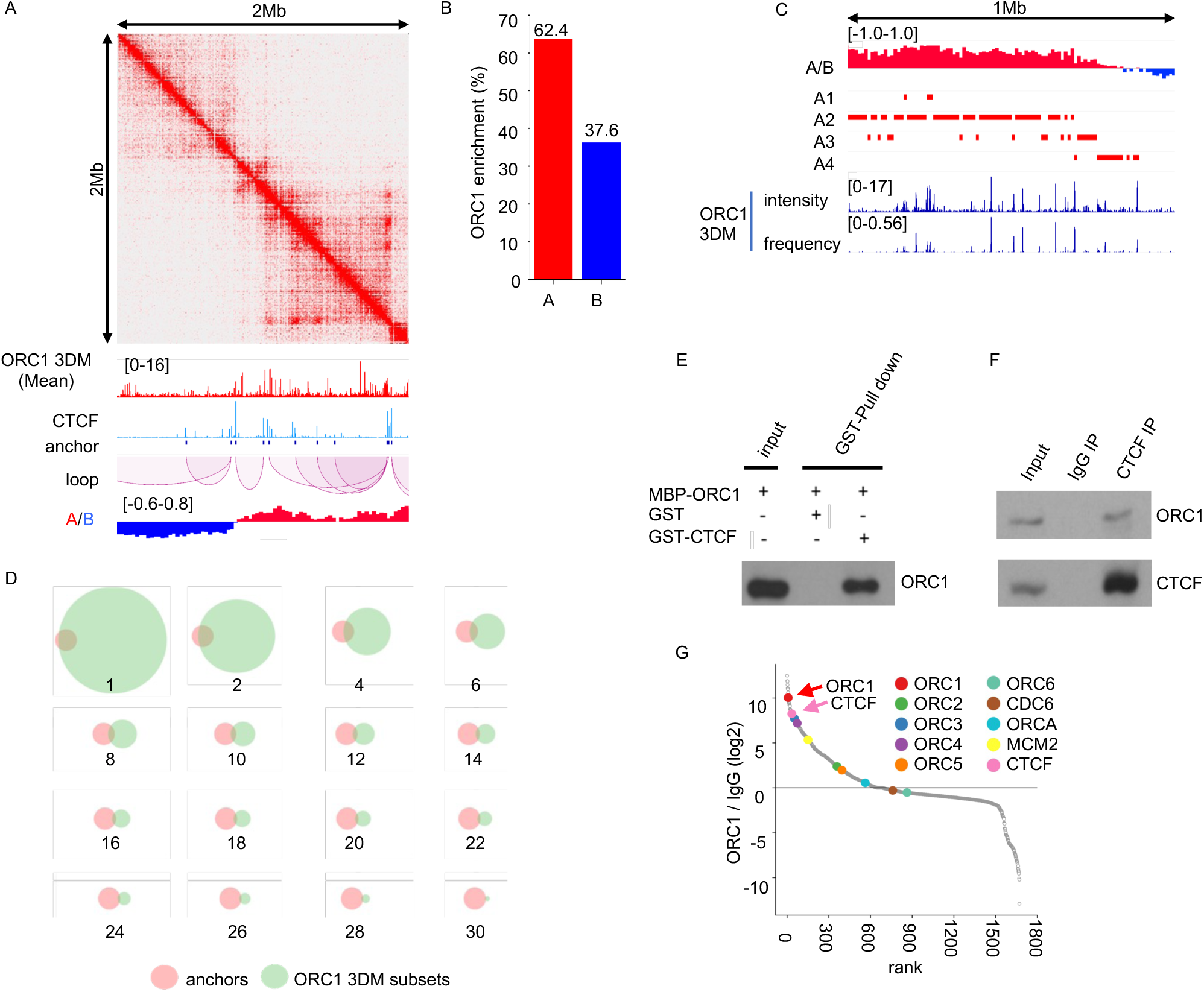
ORC1 3DM correlates with higher-order chromatin 3D structures. (A) A Hi-C map from U2OS cells for the locus chromosome chr2:117,000,000-119,000,000. IGV tracks: ORC1 3DM (mean intensity); CTCF ChIP-exo; loop anchors; loops; A/B compartments (resolution: 25kb). (B) Distribution of ORC1-binding sites in A and B compartments. (C) IGV tracks of A compartments, sub-compartments and ORC1 3DM represented by mean intensity and frequency. chr3:57,800,000-58,800,000. (D) Overlaps between loop anchors and subsets of ORC1 3DM . Subsets of ORC1–binding sites are selected with frequencies indicated by numbers below each Venn diagram. For example, “1” represents a subset of ORC1-binding sites present in 1 out of 30 libraries. (E) GST antibody pull-down assay using GST and GST-CTCF as baits and MBP-ORC1 as target followed by Western blot. (F) Immunoprecipitations (IP) for ORC1 and CTCF from U2OS cells followed by Western blot. (G) Ranked protein enrichment in ORC1 co-immunoprecipitates from U2OS cells identified by mass spectrometry. ORC1∼6, ORCA, CDC6, MCM2, and CTCF are highlighted.

A subset of ORC1 peaks overlapped loop anchors (**Fig. 3A and 3D**), raising the possibility that chromatin loops contribute to the spatial organization of a subset of replication licensing or, conversely, that ORC1 may help stabilize specific loop configurations. Because many loop anchors coincide with CTCF binding sites, we compared ORC1 3DM with CTCF ChIP-exo and observed partial overlap between them (**Fig. S3G**). Biochemical assays confirmed that CTCF and ORC1 interact directly since purified GST-CTCF pulled down MBP-ORC1 in vitro (**Fig. 3E**). Immunoprecipitation with anti-CTCF antibodies demonstrated that a subset of ORC1 bound to CTCF in vivo (**Fig. 3F**). Consistent with these observations, ORC1 co-immunoprecipitation followed by mass spectrometry identified CTCF among the highly enriched proteins. The co-immunoprecipitate also contained multiple known ORC-associated proteins, including ORC2–5 and MCM2, supporting the specificity and biological relevance of the ORC1 interactome (**Fig. 3G**). Surprisingly, canonical cohesin components, including RAD21, SMC1A, and SMC3, were not similarly enriched (**Fig. S3H**), raising the possibility that the ORC1–CTCF association may occur through a cohesin-independent mode. Taken together, these findings show that ORC1-binding sites preferentially occupy the most interactive layers within A compartments and frequently coincide with 3D structural landmarks. The physical association between ORC1 and CTCF, coupled with the lack of enrichment of canonical cohesin components in ORC1 immunoprecipitates, suggests a previously unrecognized connection between ORC1, replication licensing, and higher-order chromatin organization that may operate independently of the canonical cohesin pathway.

## ORC1 3DM in the Replication Timing Landscape

To investigate how ORC1-binding sites identified in the ORC1 3DM relate to genome-wide replication initiation, we generated an 8-fraction Repli-seq dataset from asynchronous U2OS cells, following the protocol described in Marchal et al. (80) and Zhao et al. (81), with modifications. Briefly, cells were pulse-labeled with BrdU for 30 min, stained with propidium iodide, and fractionated by flow cytometry based on DNA content into eight cell-cycle fractions (P1∼P8), with P2∼P7 covering S phase (**Fig. 4A**). DNA libraries of BrdU-containing DNA were prepared for each fraction and sequenced (**Fig. 4B**). We constructed a replication-timing profile (RTP) using our newly developed Replication Timing Profiler (RTProfiler), which enables segmentation of S phase into interpretable replication-timing states at multiple resolutions.

**Fig. 4.**
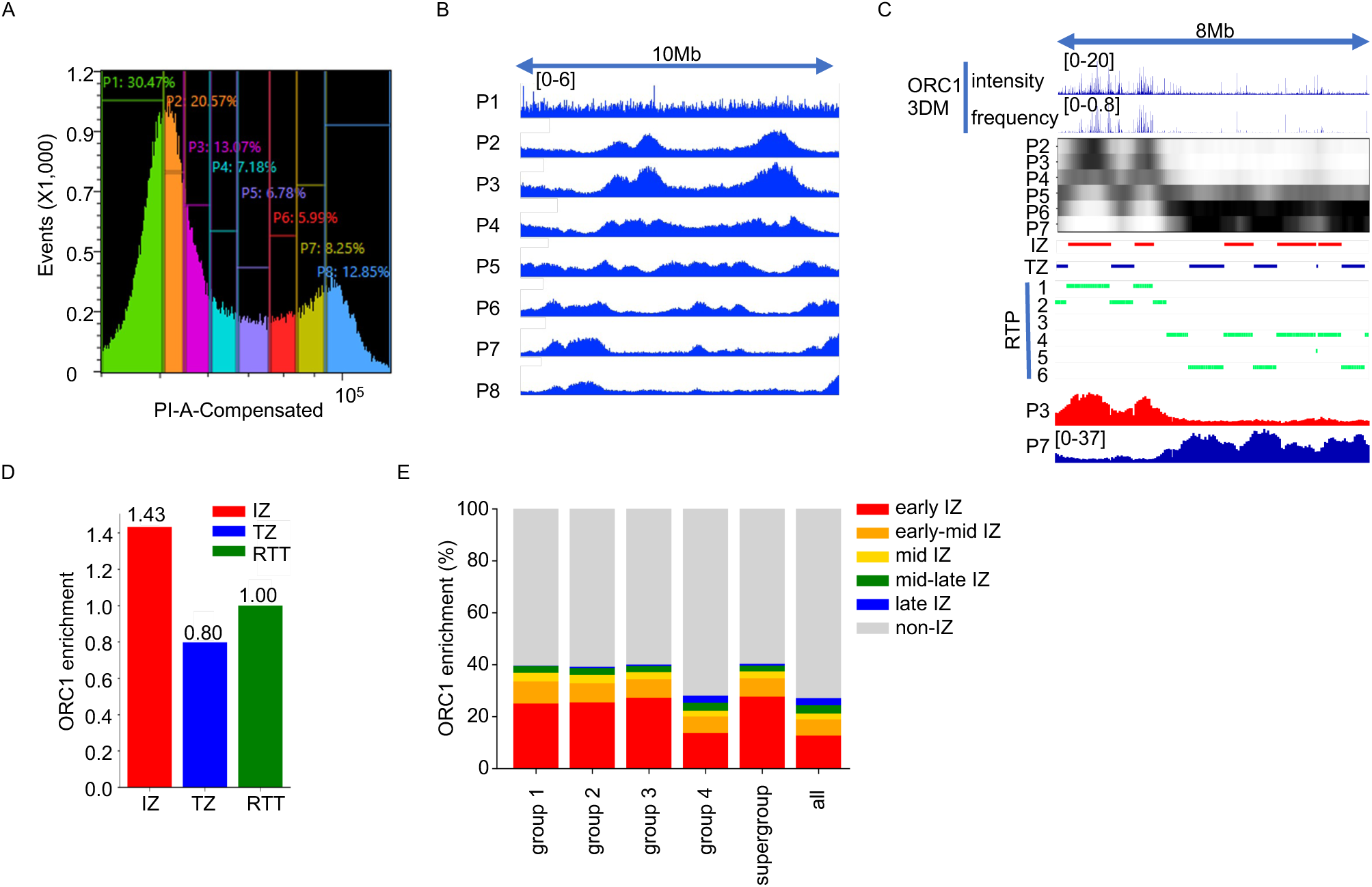
ORC1 3DM and DNA replication timing. (A) Asynchronous U2OS cells were stained with propidium iodide and sorted into eight cell-cycle fractions (P1–P8) by flow cytometry. (B) Repli-Seq IGV tracks showing coverage of the eight fractions across chr2:39,000,000–49,000,000. (C) Overlay of the ORC1 3DM with the replication-timing profile (RTP, k=6) across chr14:22,000,000–30,000,000. Track 1: ORC1 3DM frequency; track 2: ORC1 3DM mean intensity; track 3: replication-timing heatmap including p2∼p7; track 4: initiation zones (IZs); track 5: termination zones (TZs); track 6: RTP: pseudo sub-s phase 1∼6; track 7: early S-phase signal (represented by the coverage of P3); track 8: late S-phase signal (represented by the coverage of P7). (D) Enrichment of ORC1-binding sites from the ORC1 3DM in IZs, TZs, and replication timing transition zones (RTTs). Enrichment values are weighted by the genomic occupancy of the different replication-timing zones and normalized to the enrichment in RTTs. (E) Distribution of different groups of ORC1-binding sites from the ORC1 3DM across IZs and non-IZ regions in different sub-S phases.

Evaluation on chromosome 2 across k = 6, 10, 15, 20, 25, and 30 yielded 6, 10, 14, 19, 20, and 20 effective replication-timing states, respectively, indicating an approach to saturation at larger k values (**Fig. S4A**). These replication states can be ordered from early to late replication; thus, S phase can be computationally fractionated into k sub-phases. We refer to these as pseudo sub-S phases to distinguish them from the “hard” sub-S phases defined by flow cytometry. This broad range of usable k values provides flexibility: coarse-grained RTPs capture global replication domains, whereas fine-grained RTPs resolve sub–S-phase structure. Next, we mapped the RTP onto the ORC1 3DM, represented by intensity and frequency. To estimate the genome-wide distribution of ORC1-binding sites across replication-timing zones, we first used a coarse-grained RTP (k = 6) and observed that ORC1-binding sites are enriched in early-replicating regions (**Fig. 4C**). Nearly half of all ORC1-binding sites (48.4%) fall within initiation zones (IZs) (**Fig. S4B**). At k = 6, ORC1 peak density was ∼43% higher in IZs and ∼20% lower in termination zones (TZs) than in replication-timing transition regions (RTTs). The RTT category comprised peaks assigned to neither IZs nor TZs. Peak densities were normalized to RTT, giving an RTT value of 1 (**Fig. 4D**). These results confirm that ORC1 preferentially marks genomic regions that initiate replication. To examine the hierarchical relationship between ORC1-binding site groups and replication timing at finer resolution, we generated an RTP using k = 20, yielding 20 pseudo sub-S phases that were grouped into early, early-mid, mid, mid-late, and late S phases (**Fig. S4C**). IZs and TZs are present across all of these sub-S phases. The temporal ordering of these phases was confirmed by our EdU-seq data (**Fig. S4D**), in which EdU signal is maximal in early IZs and progressively decreases across later IZ states. We quantified the distribution of each ORC1 3DM group across IZs and non-IZs within these refined sub-S phases (**Fig. 4E**). All ORC1 groups were enriched in early IZs, consistent with the expected early firing behavior of many replication origins. However, group 4 exhibited a distinct pattern: while groups 1∼3 showed few ORC1 sites in late IZs, group 4 displayed a substantial fraction of peaks in both late IZs and late TZs. In TZs, groups 1–3 showed a modest total overlap of 10.8–12.3%, whereas group 4 showed approximately 20% overlap, with a notable accumulation in late TZs (**Fig. S4E**). At this higher resolution (k = 20), a majority of ORC1-binding sites no longer fall within IZs. This behavior is expected: as resolution increases, the fractional genomic coverage of IZs decreases, reducing the probability of direct overlap.

Consistent with this, when using k = 6, the fraction of ORC1-binding sites within IZs rises to ∼70% for the ORC1 supergroup (**Fig. S4F and S4G**). Collectively, these results support the prevailing model that only a subset of ORC1-binding sites fire in a given S phase (1, 15), and that distinct ORC1 subgroups exhibit characteristic timing behaviors across the S-phase continuum.

## Integrated analysis of ORC1 3DM with replication initiation landscapes and chromatin context

Numerous genome-wide DNA replication initiation landscapes generated using SNS-seq, ini-seq2, DNAscent, and OK-seq are publicly available. To compare these datasets with the ORC1 3DM within a broader chromatin context, we integrated replication initiation landscapes with a wide range of genomic features and generated a comprehensive heatmap summarizing their relationships (**Fig. 5**). For simplicity, we first collapsed Fig. 2E, which contains one genome occupation column and 30 columns corresponding to subsets of ORC1-binding sites. Based on Fig. 2E, subsets within each group show highly similar chromatin state distributions. Therefore, we used one representative subset for each group: “eq_30” for group 1, “eq_26” for group 2, “eq_15” for group 3, and “eq_3” for group 4. Notably, unlike Fig. 2E where data were normalized column-wise to compare distributions within subsets, here we used raw enrichment values for these four representative subsets to allow direct comparison between groups. This reduced heatmap serves as a structural framework. Onto this framework, we integrated additional datasets, including ini-seq2 (37), DNAscent (82), SNS-seq (83), and OK-seq (4) to represent replication initiation landscapes; Hi-C (this study) and CTCF ChIP-exo (this study) to represent 3D chromatin structure; ATAC-seq (84) and DNase-seq (85) to represent chromatin accessibility; and BG4 CUT&TAG (86) to represent G-quadruplex structures. All in-house datasets, including ORC1 3DM, CTCF ChIP-exo, and Hi-C, were generated in U2OS cells. The public ATAC-seq, DNase-seq, and BG4 CUT&Tag datasets were also derived from U2OS cells. As shown in Fig. 2E and recapitulated here, ORC1-binding sites in the ORC1 3DM dataset are primarily enriched in two types of chromatin regions: the supergroup (groups 1∼3; states 21∼23) and group 4 (states 2∼4). To simplify subsequent comparisons, we use “supergroup region” and “group 4 region” as the backbone for interpreting relationships between ORC1-binding sites, replication landscapes, and other genomic features. Ini-seq2 focuses on early S phase and includes two time points: 15 minutes and 3 hours, representing extremely early and early firing origins, respectively. Both time points are mainly enriched in chromatin states 21 and 22, while the 3-hour dataset also shows enrichment in states 23 and 17∼20. Overall, initiation sites mapped by ini-seq2 were concentrated in the supergroup region, suggesting that ORC1-binding sites in this region tended to fire early. DNAscent captures replication events across the entire S phase in asynchronous cells, revealing replication activity genome wide. Although the signal was broadly distributed, relatively higher enrichment was still observed in the supergroup region, suggesting that initiation events in this region are more conserved across cell populations. The SNS-seq dataset used here is an integrated resource compiled from 22 libraries across six different cell types. Like DNAscent, SNS-seq captures replication events throughout S phase and therefore shows a broad genomic distribution. Nevertheless, the supergroup region still exhibited relatively higher enrichment compared to other genomic regions. The OK-seq dataset used here was derived from HeLa cells and captures replication events throughout S phase. The initiation zones (IZs) defined in this study could be divided into four types of regions with distinct chromatin states: states 1∼10 (region 1), 15∼19 (region 2), 21∼23 (region 3), and 24∼25 (region 4). Among these, region 3 shows the highest enrichment and corresponds to the supergroup region. Other IZ regions overlap with the group 4 region. In contrast, non-IZ regions (TZ, HFZ, and LFZ) display lower enrichment in the supergroup region compared to the genomic background. These results further support that the supergroup region harbors more conserved initiation events. Although the relationship between ORC1 3DM and Hi-C data was analyzed in Fig. 3, here we examine this relationship from a chromatin state perspective. At the compartment level, A/B compartments show complementary chromatin state patterns. A compartments are enriched in states 2∼3, 6∼8, 16∼19, and 21∼23, whereas B compartments are enriched in states 12∼14 and 20. Both the supergroup and group 4 regions fall within A compartments. However, certain A compartment regions (states 6∼8 and 16∼19) are relatively depleted of ORC1-binding sites. At finer-scale structures such as TAD boundaries and loop anchors, the chromatin state distribution more closely resembles that of ORC1-binding sites, with stronger enrichment in the supergroup and group 4 regions. Thus, from compartments to boundaries to anchors, increasing structural resolution corresponds to increasing similarity to the ORC1-binding pattern. CTCF ChIP-exo data largely recapitulates the pattern observed for Hi-C anchors, suggesting potential functional interactions between CTCF and ORC1 at these sites. ATAC-seq, DNase-seq, and BG4 CUT&Tag datasets display chromatin state distributions highly similar to ORC1 3DM, reinforcing that ORC1-binding sites are enriched in open chromatin regions and that G-quadruplex structures may contribute to ORC1 binding or to replication progression surrounding initiation sites. Collectively, this heatmap integrates chromatin states, 3D structures, chromatin accessibility, replication initiation landscapes, and ORC1-binding profiles into a unified framework, providing a foundation for deeper mechanistic analysis.

**Fig. 5.**
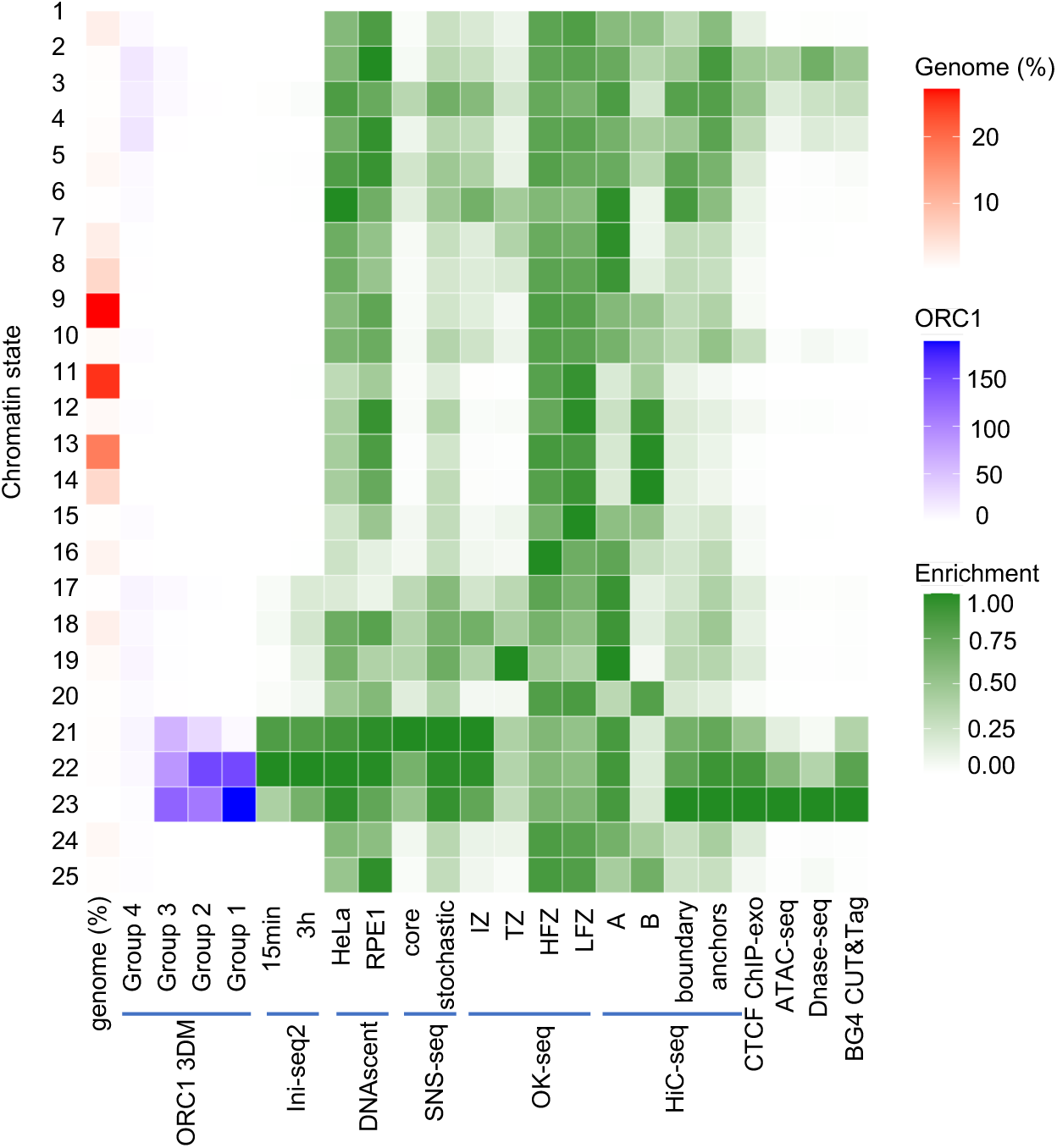
Heatmap integrating ORC1 3DM with chromatin states, replication initiation landscapes, and genome features. Genome (%) indicates the genome coverage of the corresponding chromatin state. ORC1 3DM: groups 1∼4 are represented by the subsets marked with an asterisk in Figure 2E. Ini-seq2 in EJ30 cells at two time-points. DNAscent in HeLa and RPE1 cells. SNS-seq: two sets of origins, core and stochastic, mapped in 22 libraries across 6 cell types. OK-seq in HeLa cells. Initiation zone (IZ), termination zone (TZ), high flat zone (HFZ), low flat zone (LFZ). Hi-C seq in U2OS cells. CTCF ChIP-exo in U2OS cells. ATAC-seq in U2OS cells. DNase-seq in U2OS cells. BG4 CUT&TAG in U2OS cells.

## Discussion

Our results demonstrate that ORC1 binding in human cells is reproducible yet intrinsically heterogeneous, and that this heterogeneity reflects meaningful biological structure rather than experimental noise. Although individual ORC1 CUT&RUN datasets show sharp enrichment and strong separation from IgG controls, quantitative comparisons across replicates reveal substantial variability in peak detection. This behavior is consistent with observations from other CUT&RUN and CUT&Tag studies and highlights a fundamental limitation of single-replicate, peak-centric representations of broadly distributed chromatin-associated proteins. Importantly, degron-mediated acute and short-term depletion of ORC1 leads to a marked loss of CUT&RUN signal without inducing overt cellular defects, confirming that the observed enrichment patterns are ORC1-dependent and biologically specific.

The partial but structured co-localization between ORC1 and other licensing components underscores the complexity of origin recognition in human cells. While ORC subunits, CDC6, ORCA, and MCM proteins broadly follow ORC1 binding patterns, differences in the magnitude of overlap suggest that origin licensing complexes, including all ORC subunits and CDC6, may not exist at all ORC1-bound sites. This heterogeneity may reflect dynamic licensing states, variable stability of ORC assemblies, or additional non-replicative functions of individual subunits. Although differences in antibody performance cannot be fully excluded, the consistent hierarchy observed across datasets argues for biologically meaningful variation in ORC complex composition or occupancy. Indeed, it is known that ORC2 and CDC6 function at centromeres and at sites of gene regulation, independent of their role in licensing the initiation of DNA replication (71, 73). The quantitative analysis of the ORC1 CUT&RUN data also resolves a long-standing issue about the lack of overlap between previous ORC ChIP-Seq datasets and the observation that different ORC subunits do not localize to the same sites (41, 56, 60). Our new data show extensive overlap between ORC subunits, CDC6, MCM subunits, and ORC-binding proteins, consistent with their joint role in establishing pre-RCs to license origins of DNA replication. A recent CUT&RUN mapping of ORC6 binding sites in the human genome (87) used SEACA for peak calling and the overlap with SNS-seq initiation sites was only 26.9%.

However, a re-analysis of their data using MACS2 shows considerable overlap with our ORC subunit data and sites of initiation of DNA replication (data not shown).

A central conceptual advance of this work is the development of the ORC1 3DM, which explicitly decouples binding intensity from reproducibility across experiments. Traditional peak-calling approaches implicitly assume that stronger signal equates to higher biological relevance; however, our data revealed that ORC1 peaks with comparable intensities can differ dramatically in detection frequency. By integrating reproducibility as an independent dimension, ORC1 3DM captures both stable and variable components of ORC1 occupancy in the genome, allowing population-level binding behavior to be quantified rather than the binding being reduced to a simple, uniform map. This framework establishes variability as an informative property of ORC1 binding and provides a basis for future correlation of binding sites with biological function.

Chromatin-state analysis of ORC1 3DM reveals a hierarchical organization of ORC1 binding that is tightly linked to epigenomic context, similar to a correlation with origin efficiency and chromatin features (37). High-frequency ORC1-binding sites preferentially occupy chromatin states enriched for H3K4me3 and H2A.Z1, features associated with open, transcriptionally permissive chromatin and high nucleosome turnover. Lower-frequency ORC1 sites, by contrast, are biased toward more compact or weakly repressive chromatin environments. These findings indicate that ORC1 is capable of associating with a broad range of chromatin contexts or states but does so with varying stability or detectability. The strong association with H2A.Z1-containing chromatin, previously observed in one ORC1 ChIP-Seq data set (58), suggests that histone variant composition may play a facilitative role in ORC1 recruitment or retention, potentially by modulating nucleosome dynamics or DNA accessibility. Moreover, H3K4me3 has been associated with ORC2 ChIP-Seq (56) and with origins of DNA replication in human and *C. elegans* cells (29, 37, 88, 89), further strengthening the link between ORC1 binding sites and sites of initiation of DNA replication.

Integration of ORC1 3DM with Hi-C data further reveals that ORC1 binding is embedded within the hierarchical organization of three-dimensional chromatin structure. While ORC1-binding sites are enriched in the A compartments overall, finer subdivision of these compartments based on interaction intensity demonstrates that ORC1 preferentially localizes to the most interactive sub-regions within A compartments. This relationship cannot be explained solely by genomic coverage and indicates that ORC1 binding is sensitive to quantitative features of chromatin folding. Regions with higher local interaction density may provide a structurally favorable environment for replication licensing, either by increasing local concentration of regulatory factors or by stabilizing ORC1 association through spatial constraints.

The association of ORC1-binding sites with loop anchors adds an additional layer of spatial organization. Chromatin loops represent stable architectural features that define long-range regulatory interactions, and their frequent coincidence with ORC1 binding suggests a functional coupling between replication licensing and 3D genome topology. This observation supports two non-exclusive models: chromatin loops may spatially coordinate replication licensing events across defined genomic distances, and/or ORC1 itself may contribute to loop formation or stabilization. The partial overlap between ORC1 and CTCF binding sites, together with direct biochemical evidence of ORC1–CTCF interaction and proteomic identification of CTCF in ORC1 immunoprecipitates, lends support to both possibilities. Interestingly, proteomic analysis further revealed strong enrichment of CTCF but not canonical cohesin components, suggesting that the ORC1–CTCF interaction may occur independently of the canonical CTCF–cohesin pathway.

One possibility is that CTCF may stall or block one of the DNA replication forks emanating from origins located at the subset of ORC-binding sites that colocalize with CTCF, potentially causing unidirectional replication from these origins. Unidirectional replication in the mammalian genome has been considered based on existing data (2). Collectively, these findings support a functional connection between ORC1, replication licensing, and higher-order chromatin organization.

Replication-timing analysis provides a functional context for these structural associations. ORC1-binding sites are strongly enriched within initiation zones at coarse temporal resolution, confirming that ORC1 occupancy marks genomic regions with high initiation potential. However, as replication timing is examined at finer resolution, many ORC1-binding sites fall outside initiation zones, consistent with the well-established principle that only a subset of licensed origins fire during any given S phase (1, 15). The distinct timing behaviors of ORC1-binding subgroups further emphasize this distinction since the high-frequency ORC1 sites preferentially associate with early initiation zones, whereas lower-frequency groups—particularly group 4— extend into late initiation and termination zones. These patterns suggest that different classes of ORC1 binding may correspond to origins with distinct firing probabilities, efficiencies, or regulatory roles.

Notably, the enrichment of group 4 ORC1-binding sites in late initiation and termination zones raises the possibility that some ORC1 associations reflect dormant origins, backup licensing sites, or ORC1 functions that are not directly coupled to replication initiation. The convergence of chromatin-state, 3D-structure, and replication-timing differences supports a model in which ORC1 binding is not a single uniform phenomenon, but instead encompasses multiple functional modes shaped by chromatin accessibility, spatial organization, and cell-cycle regulation.

The integrated heatmap further situates these subgroup-specific behaviors within a broader replication and chromatin framework. Across multiple independent initiation-mapping technologies, initiation events show preferential enrichment in the supergroup region (states 21∼23), where high-frequency ORC1-binding sites are concentrated, indicating that this chromatin environment represents a recurrent and conserved context for origin firing. Although replication activity is detectable genome-wide across S phase, early and population-conserved initiation events are disproportionately associated with this region. At the level of 3D genome organization, higher-resolution structures such as TAD boundaries and loop anchors exhibit chromatin state distributions that more closely parallel ORC1-binding patterns than do bulk compartments, reinforcing the link between replication licensing and local chromatin topology (78). The concordant enrichment of ORC1-binding sites with open chromatin features and G-quadruplex structures further supports a model in which epigenomic state, DNA structural features, and spatial organization collectively constrain the landscape of potential initiation sites.

Overall, these findings support a probabilistic and hierarchical view of replication licensing in human cells. In this view, ORC1 binding defines a structured and context-dependent landscape of potential initiation sites constrained by chromatin state and three-dimensional genome architecture, while replication firing represents a selective process acting on this landscape, with distinct ORC1 subgroups corresponding to origins of differing firing probability, efficiency, or regulatory function.

## MATERIALS AND METHODS

Experiments were performed in human U2OS cells. Endogenous ORC1 was tagged with FKBP12^F36V^ using CRISPR/Cas9 to enable dTAG-mediated depletion. ORC1 binding was profiled by CUT&RUN, with IgG and ORC1-depletion controls used to assess specificity. Additional replication proteins and histone modifications were profiled by CUT&RUN or CUT&Tag. Sequencing reads were aligned to the human hg38 reference genome. ORC1 3DM was constructed from 30 CUT&RUN datasets meeting quality thresholds of more than 5,000 peaks and FRiP ≥0.1. MACS2 peaks were mapped to 50-bp genomic bins, excluding ENCODE blacklist regions. Signals were normalized to a reference dataset by histogram matching and scaling to the reference mean and standard deviation. Each bin was characterized by its genomic position, peak-detection frequency across datasets, and normalized signal intensity; signal means and standard deviations were calculated from nonzero observations. ChromHMM defined 25 chromatin states from ten histone modifications at 200-bp resolution. RTProfiler used K-means-initialized Gaussian mixture modeling to infer ordered replication-timing states and identify putative replication initiation and termination zones. ORC1 binding was integrated with chromatin states, chromosome organization, replication timing, and published initiation datasets. Protein interactions were examined by GST pull-down, coimmunoprecipitation, and mass spectrometry. Chromosome organization was assessed by in situ Hi-C and CTCF ChIP-exo, and replication timing was measured by eight-fraction Repli-seq and EdU-seq.

Detailed experimental protocols, reagents, and computational procedures are provided in the SI Appendix.

## AI assistance

OpenAI Codex (GPT-6 Astra) was used to assist with coding, data analysis, manuscript editing, preparation of the Materials and Methods summary, and reference checking and formatting. The authors reviewed and verified the resulting text and references and take responsibility for the final manuscript.

## Data and materials availability

Plasmids and U2OS ORC1-FKBP12^F36V^ cell lines generated in this study are available from the lead contact without restriction. All data generated in this study have been deposited to the Gene Expression Omnibus with the open accession GSE351020 and will be made publicly available upon publication. The scripts used for data processing will be made publicly available on GitHub (https://github.com/meo2010/ORC1-3DM) upon publication under the MIT License.

## ACKNOWLEDGMENTS

We thank members of the Stillman lab for critical discussions. This work was funded by NIH grants CA13106 and GM45436, and the Goldring Family Foundation. The use of shared research resources at Cold Spring Harbor Laboratory was facilitated by a Cancer Center Grant (CA13106).

## AUTHOR CONTRIBUTIONS

M.O. and B.S. conceived and designed the experiments and analyzed the data. M.O. performed the experiments. M.O. and B.S. wrote and edited the paper and designed and drew the figures.

## DECLARATION OF INTERESTS

The authors declare no competing interests.

## Notes

### Competing Interest Statement

The authors have declared no competing interest.

