## Supplementary material for "Specification of Origins of DNA Replication Within Distinct Chromatin States in Human Cells": Suplemental Material and Data

### Materials and Methods

Detailed experimental procedures and analysis parameters are provided below. References are numbered independently within this Supporting Information.

#### Cell lines and culture conditions

U2OS cells were obtained from the Cold Spring Harbor Laboratory cell culture collection and cultured in DMEM containing high glucose (GIBCO) supplemented with 10% inactivated fetal calf serum. This cell line was tested negative for mycoplasma contamination. ORC1-FKBP12$^{F36V}$ U2OS cell lines were cultured in the same conditions as U2OS cells.

#### Plasmid construction

To generate the CRISPR gRNA/Cas9 construct (pX330-1X2-ORC1-5-gRNA) for creation of the ORC1-FKBP12$^{F36V}$ U2OS cell line, a guide RNA (gRNA) targeting the 5′ end of the human ORC1 gene (top strand: 5’-CACCGgtagtgtgccatggcttctg-3’; bottom strand: 5’-AAACcagaagccatggcacactacC-3’) was cloned into the CRISPR/Cas9 expression vector pX330-1X2 (Addgene, #58766) according to a standard protocol (1). To generate the donor construct, a 1.4 kb genomic fragment encompassing the ORC1 start codon at its center was amplified from U2OS genomic DNA using the following primers: forward, 5’-gggcccccctcgagATTGCAGTCTGGCTTTTCCGG-3’; reverse, 5’-ATCGATACCGTCGACGGGAAACCAATTCGCCCAG-3’. The PCR product was inserted into the pBluescript backbone by In-Fusion cloning to generate pBluescript-ORC1-5. The BSD-P2A-2×HA-FKBP12$^{F36V}$ cassette was amplified from pCRIS-PITChv2-BSD-dTAG-BRD4 (Addgene, #91792) using primers: forward, 5’-TCCACAGAAGCCATGATGGCCAAGCCTTTGTCTCAAG-3’; reverse, 5’-TGTGGGGTAGTGTGCGCCAGATCCGCCGCCACC-3’. The amplified cassette was inserted into pBluescript-ORC1-5 by In-Fusion cloning to generate the final donor plasmid, pBluescript-ORC1-5-BSD-P2A-2×HA-FKBP12$^{F36V}$. The resulting construct contained 787 bp and 607 bp left and right homology arms, respectively.

For protein expression constructs, human CTCF cDNA from the FLEXgene collection was amplified using primers: forward, 5’-TTCCAGGGGCCCCTGGGATCCATGGAAGGTGATGCAGTC-3’; reverse, 5’-GTGATGATGGTGATGGTCgacCCGGTCCATCATGCTGAG-3’, and cloned into the pGEX-6P-1 vector by In-Fusion cloning to generate GST-hCTCF-6×HIS. The MBP-hORC1-6×HIS construct was obtained from our laboratory collection.

#### Genome editing

The endogenous ORC1 locus in U2OS cells was modified by CRISPR/Cas9-mediated homologous recombination to introduce a BSD-P2A-2×HA-FKBP12$^{F36V}$ cassette at the N-terminus of ORC1. On day 1, 0.2 million wild-type U2OS cells were seeded per well in a 12-well plate. On day 2 (~70%

confluency), cells were co-transfected with pX330-1X2-ORC1-5-gRNA and pBluescript-ORC1-5-BSD-P2A-2×HA-FKBP12$^{F36V}$ using Lipofectamine 2000 according to the manufacturer's instructions. On day 7, cells were passaged to a 10-cm dish. On day 8, selection was initiated with 10 μg/ml blasticidin (Thermo Fisher, J67216-XF). Medium was replaced and selection maintained every 4 days until single colonies grew to sufficient size for picking. On day 22, individual colonies were isolated and expanded separately under continuous blasticidin selection until adequate cell numbers were obtained for analysis.

Genomic DNA was prepared from each clone. The 5′ end of the ORC1 locus was amplified by PCR using the following primers: forward, 5′-ATTGCAGTCTGGCTTTTCCG-3′; reverse, 5′-GGCCTGCCAACCCATGAATA-3′. The expected PCR product sizes were 0.8 kb for the wild-type allele and 1.7 kb for the targeted allele. Clones displaying only the 1.7 kb band were considered homozygous knock-in candidates.

Protein expression was validated by western blot using anti-ORC1 and anti-HA antibodies. Correctly targeted clones showed a single band at approximately 120 kDa corresponding to HA-FKBP12$^{F36V}$-ORC1, with loss of the wild-type ~100 kDa band. For functional validation, cells were treated with dTAGv1 for 1, 2, and 4 hours. HA-FKBP12$^{F36V}$-ORC1 levels showed time-dependent depletion upon treatment. Three independent clones passed genotyping, protein expression, and degradation validation, and the clone 1 (named as ORC1-FKBP12$^{F36V}$ U2OS for simplicity) was selected for subsequent experiments (Fig. S1C).

**Expression and purification of recombinant proteins**

GST-6xHis, GST-CTCF-6×His and MBP-ORC1-6×His fusion proteins were expressed in E. coli BL21 cells transformed with the corresponding expression constructs. Overnight starter cultures were expanded into 500 ml LB medium and grown at 37 °C to mid-log phase. Protein expression was induced with 0.5 mM IPTG, and cultures were incubated at 16 °C for approximately 9 h before harvesting. Cells were collected by centrifugation and stored at −80 °C until purification. Pellets were thawed on ice and resuspended in lysis buffer (50 mM Tris-HCl pH 7.5, 150 mM NaCl, 0.02% NP-40, 5 mM $MgCl_2$, 5 mM benzamidine-HCl, 1 mM PMSF, 1 mM DTT, 10% glycerol) supplemented with lysozyme. After incubation on ice, lysates were sonicated and clarified by centrifugation at 14,000 rpm for 45 min at 4 °C. Supernatants were supplemented with 5 mM imidazole and incubated with Ni-NTA agarose (QIAGEN, 30210) for 3 h at 4 °C with rotation. The resin was washed with high-salt wash buffer (50 mM Tris-HCl pH 7.5, 500 mM NaCl, 0.02% NP-40, 5 mM $MgCl_2$, 1 mM DTT) followed by lysis buffer, and bound proteins were eluted with 0.25 M imidazole in lysis buffer. For tandem affinity purification, Ni-NTA eluates were incubated with amylose resin (NEB, E8021S) (MBP-ORC1-6×His) or glutathione resin (Thermo scientific, 16100) (GST-CTCF-6×His) equilibrated in lysis buffer. After washing, proteins were eluted with lysis buffer containing 20 mM maltose (MBP-ORC1-6×His) or 20 mM reduced glutathione (GST-CTCF-6×His). Eluted fractions were analyzed by SDS-PAGE and Coomassie staining. Purified proteins were dialyzed overnight against 1 L of 50 mM Tris-HCl (pH 7.5) at 4 °C, followed by a second dialysis in fresh buffer for 4 h. Proteins were then adjusted to storage conditions by supplementing glycerol to a final concentration of 50% (v/v) in 50 mM Tris-HCl (pH 7.5). Samples were concentrated using centrifugal filter units (Amicon Ultra, 3 kDa cutoff) and aliquoted prior to storage at −80 °C.

pAG/MNase (Addgene, 123461), used in CUT&RUN assays, was a gift from Dr. Steven Henikoff. JM101 cells were transformed with the pAG/MNase plasmid. Overnight starter cultures were expanded into 3 L LB medium and grown at 37 °C to mid-log phase. Protein expression was induced with 2 mM IPTG, and cultures were incubated at 16 °C for approximately 9 h before harvesting. Cells were collected by centrifugation and stored at −80 °C until purification. Because the AG-MNase fusion protein contains a C-terminal 6×His tag, it was purified using the same Ni-NTA protocol described above up to the elution step. Eluted protein was further purified by gel filtration chromatography using 20 mM Tris-HCl (pH 7.5) to remove residual imidazole and contaminating proteins. The purified protein was adjusted to storage conditions by adding glycerol to a final concentration of 50% (v/v) in 50 mM Tris-HCl (pH 7.5), aliquoted, and stored at −80 °C.

**Pull down assay**

GST pull-down assays were carried out using purified recombinant proteins. Anti-GST antibody was first incubated with Dynabeads Protein G (Invitrogen, 10004D) for 15 minutes at room temperature in binding buffer (25mM Tris-HCl pH 7.5, 75 mM KCl, 0.1% NP-40, 0.1mM EDTA, 5mM MgCl2, 10% glycerol, 1mM DTT, cOmplete™ EDTA-free Protease Inhibitor Cocktail). The beads were washed three times with binding buffer and resuspended for subsequent incubation. Purified GST-hCTCF or GST control protein was added to the antibody-bound beads and incubated overnight at 4°C with rotation to allow immobilization of the GST fusion proteins. After washing three times to remove unbound material, the beads were incubated with purified MBP-ORC1 in binding buffer at 4°C for 2 hours with rotation. The beads were then washed three additional times with binding buffer. Bound proteins were eluted by heating in 1× Laemmli sample buffer (62.5 mM Tris-HCl pH 6.8, 2% SDS, 2.5% 2-mercaptoethanol, 10% glycerol, and 0.005% bromophenol blue) at 85°C for 5 minutes. An aliquot of MBP-ORC1 was reserved as input control and mixed with wash buffer and 2× Laemmli sample buffer (125 mM Tris-HCl pH 6.8, 4% SDS, 5% 2-mercaptoethanol, 20% glycerol, and 0.01% bromophenol blue), followed by heating at 85°C for 5 minutes. Eluted samples and input controls were separated by 8% SDS–PAGE. Gels were either stained with Coomassie Brilliant Blue for loading control or transferred to membranes for immunoblot analysis using the anti-ORC1 antibody (Bethyl, A301-892A).

**Co-immunoprecipitation**

CTCF co-immunoprecipitation was performed using Dynabeads Protein G. Briefly, 40 μl of beads per reaction were washed three times with wash buffer (20 mM Tris-HCl pH 7.5, 100 mM NaCl, 5 mM $MgCl_2$, 0.15% NP-40, 1 mM DTT, cOmplete™ EDTA-free Protease Inhibitor Cocktail). Beads were incubated with rabbit IgG (negative control) or anti-CTCF antibody (Cell Signaling Technology, 3418S) and rotated at 4°C for antibody coupling. 1 × $10^7$ U2OS cells were lysed in 500 μl lysis buffer (20 mM Tris-HCl pH 7.5, 250 mM NaCl, 5 mM $MgCl_2$, 0.1 mM EDTA, 0.3% NP-40, 1 mM DTT, 2 mM $CaCl_2$, 0.25U/ul Benzonase, 20uM MG-132, cOmplete™ EDTA-free Protease Inhibitor Cocktail, PhosSTOP) for 30 min on ice with intermittent mixing. Lysates were diluted 1:1 with dilution buffer (20 mM Tris-HCl pH 7.5, 5 mM $MgCl_2$, 2 mM EGTA, 1 mM DTT, 20uM MG132, cOmplete™ EDTA-free Protease Inhibitor Cocktail, PhosSTOP) and clarified by centrifugation at 16,000 × g for 15 min at 4°C. Cleared extracts (300 μl per reaction) were incubated with antibody-coupled beads overnight at 4°C with rotation. Beads were then washed three times with wash buffer. Immunocomplexes were eluted by addition of 1× Laemmli sample buffer and heating at 85°C for 5 min. Input samples (50 μl lysate) were mixed with 4× Laemmli sample buffer and processed in parallel. Eluted samples and input controls were separated by 8% SDS–PAGE and transferred to membranes for immunoblot analysis using the anti-ORC1 antibody (Bethyl, A301-892A) and anti-CTCF antibody (Cell Signaling Technology, 3418S).

**Co-immunoprecipitation followed by mass spectrometry**

U2OS cell extracts were prepared by hypotonic lysis (10 mM Tris-HCl pH 7.5, 1 mM KCl, 1.5 mM $MgCl_2$, 1 mM DTT, 0.25U/ul Benzonase, 20uM MG132, PhosSTOP, cOmplete™ EDTA-free Protease Inhibitor Cocktail) followed by Dounce homogenization. Lysates were adjusted to 150 mM NaCl and clarified by centrifugation. Protein A/G magnetic beads were coupled with either anti-ORC1 antibody or rabbit IgG and incubated with clarified lysates at 4°C. Following extensive washing with PBS, bound proteins were retained on the beads for proteomic analysis. ORC1 and IgG co-immunoprecipitate samples were submitted to the Cold Spring Harbor Laboratory Proteomics Core Facility for liquid chromatography–tandem mass spectrometry (LC–MS/MS) analysis. Proteins were identified by database searching against the UniProt human proteome, and peptide-spectrum matches were filtered to a false discovery rate of <1%. Label-free quantification was used to estimate protein abundance across samples. Protein enrichment was calculated by comparing label-free quantification intensities between ORC1 and IgG immunoprecipitates. For Fig. 3G, proteins were ranked according to their enrichment in ORC1 co-immunoprecipitates relative to the IgG control and displayed as a ranked enrichment plot. For Fig. S3H, enrichment of CTCF and cohesin-related proteins was calculated as the log2-transformed ratio of ORC1 co-IP to IgG co-IP signal, where positive values indicate enrichment in ORC1 co-immunoprecipitates and negative values indicate depletion relative to the control.

**Drug treatments**

To induce degradation of endogenous ORC1, ORC1-FKBP12$^{F36V}$ U2OS cells were treated with 100 nM dTAG v-1 (Tocris, #6914) for the indicated times. Control cells received an equivalent volume of DMSO. Following treatment, cells were harvested by trypsinization and processed for either western blotting or CUT&RUN.

**Nuclei Preparation for CUT&RUN and CUT&Tag**

U2OS cells (~10 million cells per 15-cm plate) were harvested by trypsinization, neutralized with growth medium, and pelleted at 600 × g for 5 min. Cell pellets were resuspended in PBS, pelleted again, and then resuspended in ice-cold NE1 buffer (20 mM HEPES-KOH pH 7.9, 10 mM KCl, 0.1% Triton X-100, 20% glycerol, 0.5 mM spermidine). After vortexing and incubation on ice for 10 min, nuclei were pelleted, resuspended in PBS, and lightly crosslinked with 0.1% formaldehyde for 2 min at room temperature. Crosslinking was quenched with 60 µl of 1.25 M glycine, and nuclei were pelleted at 4°C. The nuclei were washed once, resuspended in wash buffer (20mM HEPES pH7.5, 150mM NaCl, 0.5mM Spermidine, cOmplete™ EDTA-free Protease Inhibitor Cocktail) supplemented with 10% DMSO, and cryopreserved at −80°C using a Mr. Frosty freezing container.

**CUT&RUN**

Two CUT&RUN protocols were employed. Histone marks (H3K4me3, H3K9ac, H3K9me3, H3K27ac, H3K27me3) CUT&RUN was performed using the ChIC/CUT&RUN Kit (EpiCypher, 14-1048) according to the manufacturer's instructions.

ORC1, ORC2 and H2A.Z1 CUT&RUN was performed as previously described (2) with minor modifications. Briefly, thawed nuclei were sequentially washed in PBS, Buffer I (20 mM HEPES pH 7.5, 150 mM NaCl, 2 mM EDTA, 0.1% BSA, 0.5 mM spermidine), and Buffer II (20 mM HEPES pH 7.5, 150 mM NaCl, 0.1% BSA, 0.5 mM spermidine), followed by blocking in HNT-Preblock buffer (20 mM HEPES pH 7.5, 100 mM NaCl, 1mM PMSF, 0.1% Tween-20, 3% BSA, 2mM EDTA, cOmplete™ EDTA-free Protease Inhibitor Cocktail), HNT-PPi buffer (20 mM HEPES pH 7.5, 100 mM NaCl, 1mM PMSF, 0.1% Tween-20, 0.1% BSA, cOmplete™ EDTA-free Protease Inhibitor Cocktail). Nuclei were resuspended at 6 × $10^6$/ml, and 1.8 × $10^6$ nuclei were used per CUT&RUN reaction. Primary antibody incubation was performed overnight at 4°C with rotation, followed by washes in HNT-Pi wash buffer (20 mM HEPES pH 7.5, 100 mM NaCl, 1mM PMSF, 0.1% Tween-20) and secondary antibody incubation for 1 h at 37°C. After additional washes, nuclei were incubated with AG-MNase fusion protein in Buffer II for 30 min at 37°C, washed, and then resuspended in Buffer II containing 2 mM $CaCl_2$. Chromatin digestion proceeded for 60 min at 4°C, after which reactions were stopped with an EDTA/EGTA master mix (final 10 mM EDTA, 2 mM EGTA). Released chromatin fragments were extracted, followed by deproteinization in DNA extraction buffer (20 mM Tris-HCl pH 8, 10 mM EDTA, 2 mM EGTA, 0.1% SDS, 300 mM NaCl, 0.17ug/ul Proteinase K, and 0.33ug/ul RNase A) and purification using PCI/CI extraction and ethanol precipitation. DNA was size-selected to ≤700 bp using AMPure XP beads and quantified with a Qubit fluorometer.

**CUT&Tag**

ORC3~6, ORCA, CDC6, MCM2~3 binding sites and some of the histone marks, i.e. H3K36me3, H3K56ac, H4K20me1 and H4K20me3 are mapped genome-wide using a modified CUT&Tag assay developed by Lu et al. (3). In brief, this modified CUT&Tag assay was performed using antibody-guided Tn5 tagmentation followed by T7-mediated linear amplification. Briefly, 50,000 nuclei per reaction were immobilized on magnetic beads and incubated with primary antibodies, followed by T7-adapter-loaded nanobody-Tn5. After washing, tagmentation was performed at 37°C for 1 h. Genomic DNA was purified using AMPURE XP beads, followed by gap filling and in vitro transcription using T7 RNA polymerase at 37°C for 16~19 h. The resulting RNA was purified and reverse-transcribed using random hexamer primers. Following RNase H treatment and second-strand synthesis, cDNA was fragmented and adapter-tagged using Tn5 transposase. Sequencing libraries were amplified by 15 cycles of PCR using Nextera index primers, purified using AMPURE XP beads, and assessed for fragment size distribution using a TapeStation analysis. The resulting libraries were sequenced on an Illumina NextSeq 2000 platform (paired-end 50bp).

**Library Preparation and Sequencing**

Sequencing libraries were prepared using the NEBNext Ultra II DNA Library Prep Kit (NEB) following the manufacturer's instructions. Libraries were purified with AMPure XP beads (0.8× ratio), eluted in 20 µl $H_2O$, and analyzed on a TapeStation (Agilent). Libraries were pooled followed by quality assessment. Pooled libraries were sequenced on an Illumina NextSeq 2000 platform using paired-end 50 cycles or 100-cycle P3 XLEAP chemistry kits.

**In Situ Hi-C**

In situ Hi-C was performed in U2OS cells as previously described (4) with modifications. Briefly, crosslinked nuclei were lysed and digested overnight with MboI. Overhangs were filled in with biotinylated nucleotides, followed by proximity ligation. Crosslinks were reversed, and DNA was purified, sonicated to 300–500 bp, and size-selected. Biotinylated DNA fragments were pulled down with streptavidin T1 beads, end-repaired, A-tailed, and ligated to indexed adapters. After titration PCR to determine the optimal cycle number, libraries were amplified using Q5 polymerase, purified with AMPure XP beads, and quality-controlled by TapeStation analysis. Libraries from four biological replicates were sequenced on an Illumina NextSeq 2000 platform (paired-end 75 bp) across two sequencing runs.

**EdU-seq**

EdU-seq was performed in U2OS cells as previously described (5, 6) with modifications. Briefly, cells were seeded in 15-cm plates (10 million cells per plate; three plates) and synchronized in G1 phase using 1 μM palbociclib for 20 h. Cells were released from G1 arrest by washing three times with PBS and then incubated with 10μM EdU in the presence of 2 mM hydroxyurea (HU) and 0.5 μg/ml aphidicolin (APH). After 3 h of labeling, cells were washed three times and released into fresh medium for 30 min prior to harvesting. Cell pellets were resuspended in PBS and fixed by addition of ice-cold methanol (final concentration 90%) at −20°C. Fixed cells were permeabilized with 0.2% Triton X-100 in PBS for 30 min at room temperature and subjected to click chemistry to conjugate biotin to incorporated EdU using a reaction mixture (85.5mM Tris-HCl pH 8.0, 4mM $CuSO_4$, 100mM sodium ascorbate, and 50uM Azide-PEG3-biotin). Following incubation for 30 min at room temperature in the dark, cells were washed and lysed in lysis buffer (10 mM Tris-HCl pH 8.0, 10 mM EDTA, 0.5% SDS, and 0.2 g/L proteinase K), and incubated at 50°C for >3 h. Genomic DNA was purified by phenol–chloroform extraction using Phase Lock Gel tubes, followed by chloroform extraction and ethanol precipitation. DNA pellets were washed with 70% ethanol, air-dried, resuspended in TE buffer (10mM Tris-HCl pH 8.0, 1mM EDTA), and quantified by Qubit fluorometry. For enrichment of nascent DNA, 15 μg of purified DNA was sonicated to 100–500 bp fragments and incubated with Dynabeads MyOne Streptavidin C1 (50 μl beads per sample). After rotating at room temperature for 15 min, beads were washed four times with high-salt binding buffer (5mM Tris-HCl pH 7.5, 1mM EDTA, 1 M NaCl, 1% Tween 20), with the final wash performed in TE buffer. Biotinylated DNA was eluted in elution buffer (10mM Tris-HCl pH 8.0, 2% β-mercaptoethanol) and quantified. Sequencing libraries were prepared using the NEBNext Ultra II DNA Library Prep Kit for Illumina (NEB, E7645L) according to the manufacturer's instructions. Libraries were amplified for 13 PCR cycles (98°C for 10 s, 65°C for 75 s), following an initial denaturation (98°C for 30 s) and final extension (65°C for 5 min). PCR products were purified using 0.9× AMPURE XP beads, eluted in 0.1× TE buffer, and quality-assessed by TapeStation analysis. Libraries were sequenced on an Illumina NextSeq 2000 platform (paired-end 75 bp, High Output mode).

**CTCF ChIP-exo**

ChIP-exo (version 5.0) was performed in U2OS cells as previously described (7) with modifications. Briefly, U2OS cells were cultured in 15-cm plates to approximately 20 million cells per plate. Cells were harvested by trypsinization, and 40 million cells (from two plates) were combined to generate one pellet. A total of six pellets (for six replicates) were prepared. Cells were crosslinked in 1% formaldehyde (freshly prepared in PBS) for 10 minutes at room temperature with rotation, and the reaction was quenched with 1.25 M glycine for 5 minutes. Pellets were washed twice with PBS, flash-frozen in liquid nitrogen, and stored at −80°C until use. Then, the remaining steps followed the protocol described in (7) except the antibodies and oligos. The antibodies used in this study were rabbit mAb IgG isotype control (Cell Signaling Technology, 3900s) and anti-CTCF antibody (Cell Signaling Technology, 3418S). The oligos used in this study were documented in Table S1. Libraries were sequenced on an Illumina NextSeq 2000 platform using 100-cycle P3 XLEAP chemistry.

### Repli-seq

Repli-seq was performed in U2OS cells as previously described (8) with modifications. Briefly, cells were cultured in 15-cm plates and pulse-labeled with 100 µM BrdU for 30 min. Following labeling, cells were harvested, washed with PBS, and fixed in 70% ethanol at −20°C. Fixed cells were resuspended in PBS containing 1% BSA and stained with propidium iodide (3.8 mM sodium citrate, 10 µg/ml propidium iodide, 0.1 µg/ml RNase A). Cells were sorted by flow cytometry into eight fractions spanning the entire cell cycle based on DNA content. Sorted cells were pelleted and lysed in SDS–proteinase K buffer (0.2 mg/ml proteinase K) at 56°C for 2 hours. Genomic DNA was purified using a silica column–based DNA purification kit and fragmented by sonication (30s on, 30s off, 20 cycles). Sequencing libraries were prepared from fragmented DNA using the NEBNext Ultra DNA Library Prep Kit according to the manufacturer's instructions. BrdU-labeled nascent DNA was enriched by immunoprecipitation. Libraries were heat-denatured at 95°C for 10 minutes, rapidly cooled on ice, and incubated with anti-BrdU antibody in IP buffer, followed by rabbit anti-mouse IgG secondary antibody. Immune complexes were pelleted, washed, and digested with proteinase K. DNA was purified using silica column–based cleanup and eluted in water. Libraries were amplified by PCR (15 cycles) using a high-fidelity DNA polymerase, size-selected with AMPure XP beads (0.85×), and quality-controlled by TapeStation analysis. Indexed libraries were pooled and sequenced on an Illumina NextSeq 2000 platform using paired-end 50-bp reads.

### Next-generation sequencing (NGS) data analysis

This study included multiple types of NGS datasets generated in this work, including CUT&RUN, CUT&Tag, ChIP-exo, Repli-seq, EdU-seq, and Hi-C. Different CUT&RUN datasets were analyzed according to their specific purposes, including conventional peak-based analysis, differential analysis between wild-type and ORC1-degron cells, and construction of the ORC1 three-dimensional binding matrix (ORC1 3DM). In addition, several publicly available datasets were incorporated into the study. FASTQ files from OK-seq and ATAC-seq studies were reprocessed, whereas processed data from ini-seq2, DNase-seq, DNAscent, SNS-seq, and BG4 CUT&Tag studies were used directly. Except where processed datasets were used directly, sequencing data were subjected to a common basic processing workflow followed by assay-specific analyses.

### Basic processing workflow

FASTQ files were trimmed to remove adapter sequences and low-quality bases using Trim Galore (--quality 15 -j 8) and evaluated using FastQC. Clean reads were aligned to the human reference genome (hg38) using Bowtie2 (--end-to-end --very-sensitive --no-mixed --no-discordant --phred33 -I 10 -X 700 -p 8 -x hg38) or BWA-MEM, depending on the assay. The resulting SAM files were converted to BAM format and sorted using samtools. Low-quality alignments, mitochondrial reads, and reads mapping to random or unplaced contigs were excluded. Unless otherwise specified, alignments with low mapping quality and unmapped reads were removed using samtools view (-b -q 30 -F 0x04). Reads mapping to chrM, random contigs, and chrUn contigs were also excluded. PCR duplicates were identified and marked using Picard MarkDuplicates ( REMOVE_DUPLICATES=false ASSUME_SORTED=true VALIDATION_STRINGENCY=LENIENT). Thus, duplicate reads were retained in the BAM files but were excluded from downstream analyses where duplicate filtering was required. When required for visualization and quantitative signal analysis, BAM files were converted to bigWig format using deepTools bamCoverage (-bs 10 --normalizeUsing RPGC --

effectiveGenomeSize 2913022398 --ignoreDuplicates --blackListFileName hg38-blacklist.v2.bed). Peak calling was performed, where appropriate, using MACS2 callpeak (-f BAMPE, -g hs, -q 0.05 --call-summits).

### CUT&RUN and CUT&Tag

CUT&RUN and CUT&Tag datasets for histone modifications, ORC subunits (ORC2~ORC6), and ORC-associated proteins, including CDC6, ORCA, MCM2, and MCM3, were processed using the basic data-analysis workflow described above.

#### CUT&RUN differential analysis for dTAG treatment

In addition to the basic processing workflow described above, residual E. coli DNA was used as a spike-in reference for normalization. Clean reads from each library were independently aligned to the E. coli reference genome (GCF_000005845.2_ASM584v2_genomic.fna) using Bowtie2 with the parameters --end-to-end --very-sensitive --no-mixed --no-discordant --phred33 -I 10 -X 700 -p 8 -x ecoli. The resulting SAM files were converted to BAM format using samtools, and low-quality alignments and PCR duplicates were filtered as described above. The number of E. coli-mapped reads in each cleaned BAM file was then determined. For each library, the spike-in scaling factor (SF) was calculated as: SF = maximum spike-in read count across all libraries / spike-in read count of the individual library. Scaled bigWig files were generated from the corresponding hg38-aligned BAM files using deepTools bamCoverage (-bs 10 --normalizeUsing None --scaleFactor SF --ignoreDuplicates --blackListFileName hg38-blacklist.v2.bed). Thus, the signal from each library was adjusted according to its relative E. coli spike-in read count. ORC1 peaks identified from DMSO-treated replicate 1 were used as a common reference set. The spike-in-scaled bigWig files were then used to generate signal matrices over these reference regions for heatmap visualization and comparison between conditions.

### ORC1 3DM

FASTQ files were processed using the basic processing workflow to generate clean BAM files. For each cleaned BAM file, the Fraction of Reads in Peaks (FRiP) was calculated to assess signal-to-noise ratio. Peaks were called using MACS2 with parameters: -f BAMPE --nolambda -g hs -q 0.05 --call-summits. Datasets with more than 5,000 peaks and a FRiP score ≥ 0.1 were selected for downstream analysis. In total, 30 high-quality biological CUT&RUN replicates were retained. Each MACS2 narrowPeak output was converted into a binary BED file by intersecting with a 50 bp tiling window (excluding ENCODE blacklist regions) using bedtools. In these files, bins overlapping peaks were labeled 1 (peak), and all other bins were labeled 0 (background). The 30 binary BED files were assembled into a 2D binary matrix, where each row represents a 50 bp genomic bin and each column represents a CUT&RUN dataset. To incorporate signal intensity (third dimension), one dataset was selected as a reference, chosen for having the highest FRiP score and a moderate number of peaks. Signal coverage was computed for each dataset in 50 bp bins using bedtools coverage. For each dataset, an elbow-plot–based method was applied to determine a dataset-specific signal cutoff, classifying bins into peak-like and background-like groups. Only peak-like bins were used for normalization. Normalization was performed in two steps: 1. Histogram matching was applied to align the shape of each dataset's peak signal distribution with that of the reference dataset. 2. Linear scaling was then used to match the mean and standard deviation to those of the reference. The normalized signal values were integrated with the corresponding binary peak matrix to construct a three-dimensional ORC1 binding matrix (ORC1 3DM). In this matrix, rows correspond to 50-bp genomic bins, columns correspond to individual CUT&RUN datasets, and each entry contains the normalized signal intensity for a given bin in a given dataset. Summary statistics were then calculated from the 3DM to derive the detection frequency (number of datasets in which a bin was detected as a peak), as well as the nonzero mean and standard deviation of signal intensity. For each genomic bin, zero-valued observations were excluded before calculating these summaries across biological replicates.

#### ChIP-exo

Using the basic processing workflow, fastq files were processed to clean BAM and bigWig files. Narrow peaks were called using MACS2 callpeak (--nolambda -q 0.05). This study generated 6 replicates for each antibody, i.e. IgG control and anti-CTCF antibody. The bigWig file of the CTCF ChIP-exo replicate 5, which has the highest peak numbers, was shown in Fig. 3A. The narrowPeak files from all the 6 CTCF ChIP-exo replicates were merged into one single bed file using “bedtools merge -d 100”. The merged bed file was used as an input dataset in Fig. S3G and Fig. 5.

#### Repli-seq data analysis using RTProfiler

To characterize replication timing at different levels of temporal granularity from six S-phase fractions, we developed RTProfiler. The workflow retains the smoothing and scaling procedures adapted from reference (9), while replacing its BIRCH-based clustering and cluster-ordering steps with K-means-initialized Gaussian mixture modeling and ordering by a replication-timing index. Reads were trimmed and aligned to hg38 as described in the basic processing workflow, with PCR duplicates removed before coverage analysis. Total and mitochondrial mapped reads were counted before exclusion of mitochondrial alignments, and a sample-specific scaling factor was calculated as total mapped reads divided by nuclear mapped reads. Nuclear read counts were multiplied by this factor. After subtraction of hg38 blacklist regions, the remaining genomic intervals were partitioned into 50-kb windows, retaining only full-length windows. Read counts were obtained using BEDTools and assembled into a matrix containing the six S-phase fractions, P2–P7; chromosome Y was excluded. The matrix was smoothed along the fraction and genomic-bin axes using a 3 × 3 Gaussian kernel with $\sigma = 1$, using a modified implementation of *make_array.py* (9). After smoothing, signals across the six fractions were scaled to sum to 100 within each nonzero genomic bin. Bins with zero signal across all fractions were excluded, removing 2,280 bins and retaining 58,350 bins for timing-state analysis.

For genome-wide state inference, each fraction was standardized across retained bins to zero mean and unit variance. K-means clustering was performed with 20 initializations and a random seed of 42, and the resulting cluster centers were used to initialize a Gaussian mixture model (GMM) with full covariance matrices and a maximum of 500 iterations. Each bin was assigned to the component with the highest posterior probability. The number of components, *k*, defined the number of inferred replication-timing states. To order these states temporally, a replication-timing index was calculated for each state:

$$\mathrm{RT}_s = \frac{\sum_{j=1}^{6} j w_{sj}}{\sum_{j=1}^{6} w_{sj}}, \quad w_{sj} = \mu_{sj} - \min_j(\mu_{sj}) + 10^{-6}. \qquad (368)$$

Here, $\mu_{sj}$ is the mean normalized signal in fraction *j* across bins assigned to state *s*, and $w_{sj}$ is obtained by subtracting the minimum of that state’s six-fraction mean profile and adding $10^{-6}$. States were ranked by increasing $RT_s$, corresponding to early-to-late replication, and designated pseudo sub-S phases. These computational states represent subdivisions of the measured fraction profiles rather than additional experimentally sorted fractions.

The choice of state number was evaluated on chromosome 2 using the same K-means-initialized GMM workflow with $k$ = 6, 10, 15, 20, 25, and 30. For each model, the 4,824 retained chromosome-2 bins were assigned to the component with the highest posterior probability. Models were evaluated using the fraction of signal variance explained, the number of bins assigned to each state, segment lengths, and separation of state-specific replication-timing indices. The effective-state count was defined as one plus the number of adjacent differences of at least 0.05 between sorted timing indices; this diagnostic did not merge or reassign states. The corresponding effective-state counts were 6, 10, 14, 19, 20, and 20, indicating an approach to saturation at larger $k$ values (Fig. S4A). We used $k$ = 6 for broad timing-domain comparisons and $k$ = 20 for finer temporal subdivision. Genome-wide models were fitted separately using the same workflow and selected $k$ values.

To identify replication-timing extrema, successive retained bins with the same ordered state were collapsed into plateaus within each chromosome. Internal plateaus with earlier timing than both neighboring plateaus were classified as putative replication initiation zones (IZs), whereas plateaus with later timing than both neighbors were classified as termination zones (TZs). Chromosome-terminal plateaus were excluded from extrema calling. At $k$ = 20, states 1–4, 5–8, 9–12, 13–16, and 17–20 were grouped as early, early-mid, mid, mid-late, and late S phase, respectively, for integration with the ORC1 3DM.

For the overall ORC1 distribution analysis at $k$ = 6, ORC1 peaks were restricted to the retained replication-timing regions. Each peak was assigned to IZ or TZ according to interval overlap, with IZ taking precedence when a peak overlapped both categories. Peaks overlapping neither category were assigned to replication-timing transition regions (RTTs). IZ and TZ intervals were intersected with the retained replication-timing regions to determine their genomic coverage. RTT coverage was defined as the retained regions outside the union of IZ and TZ intervals. ORC1 peak counts were divided by the corresponding zone lengths, and the resulting peak densities were normalized to the RTT peak density:

$$E_z = \frac{\frac{N_z}{L_z}}{\frac{N_{\mathrm{RTT}}}{L_{\mathrm{RTT}}}},$$

where $N_z$ is the number of assigned ORC1 peaks and $L_z$ is the genomic coverage of category $z$. RTT length was calculated by subtracting the combined IZ and TZ coverage from the total retained genomic coverage. Normalization to RTT gives $E_{\mathrm{RTT}} = 1$.

For Figs. 4E and S4E–F, overlap percentages were calculated using all merged intervals in each ORC1 group as the denominator, including mixed and unassigned intervals. These percentages were not normalized to zone genomic coverage.

**EdU-seq**

EdU-seq datasets were processed using the basic workflow. Deduplicated BAM files were used to compare EdU signal across the inferred replication-timing categories in Fig. S4D. Because labeling followed release from G1 in hydroxyurea and aphidicolin, this analysis provides complementary evidence of early-replication association.

**Hi-C**

Four biologically independent Hi-C replicates were generated in this study and initially processed separately. FASTQ files from each replicate were aligned to the human reference genome (hg38) using BWA-MEM (-t 8 -SP). The resulting alignments were processed using pairtools. Read pairs were parsed against the hg38 main-chromosome sizes file using pairtools parse (--drop-sam --drop-seq --output-stats --assembly hg38 --no-flip --add-columns mapq --walks-policy mask) and subsequently sorted using pairtools sort. PCR duplicates were identified using pairtools dedup (--max-mismatch 3 --mark-dups). Duplicate and unmapped read pairs were removed, and only non-duplicate pairs with mapping quality scores ≥30 for both reads were retained. Biological replicate reproducibility was assessed using HiCRep v0.2.6 on 250-kb contact matrices generated from the four replicates independently subsampled to 20 million contacts each. The high reproducibility among replicates supported merging the filtered contact pairs to maximize sequencing depth for subsequent analyses. The merged contact pairs were converted into a Cooler contact matrix at 500-bp resolution using cooler cload pairs. The contact matrix was balanced using iterative correction with cooler balance (-p 8 -c 10000), and a multi-resolution .mcool file was generated using cooler zoomify. A/B compartment profiles were calculated from the pooled Hi-C contact matrix using cooltools eigs-cis. Eigenvectors were phased using GC-content tracks with the --phasing-track option, and the first two eigenvectors were calculated using --n-eigs 2. BigWig compartment tracks were generated using the --bigWig option. Compartment profiles at 25-kb resolution were used to visualize A/B compartment organization alongside ORC1 3DM intensity in Fig. 3A. For higher-resolution analysis, compartment eigenvectors were calculated at 10-kb resolution from the pooled .mcool matrix using the same procedure. To subdivide A and B compartments, separate four-component Gaussian mixture models were fitted to positive and negative GC-phased E1 values, pooling genomic bins across chromosomes. Groups were ranked by the mean E1 value of their assigned bins, with A1–A4 ordered from the most positive to the least positive and B1–B4 from the most negative to the least negative. Directly adjacent bins on the same chromosome assigned to the same group were merged into domains. The resulting A1–A4 and B1–B4 E1-defined sub-compartment states were used to analyze the distribution of ORC1 peaks. TAD-associated insulation profiles were calculated from the pooled 10-kb Hi-C contact matrix using cooltools insulation with a 200-kb insulation window. Chromosome arms were analyzed separately to avoid spanning centromeric regions, and TAD boundaries were identified from local minima in the insulation profile using the Li thresholding method implemented in cooltools. For downstream integration analyses, boundary regions were selected with finite boundary strength >0.15, excluding bins flagged as bad, yielding 5,983 intervals. Chromatin loops were identified from the pooled 10-kb Hi-C contact matrix using cooltools dots. Expected cis contact frequencies were first calculated using cooltools expected-cis. Loop calling was performed with a maximum genomic separation of 10 Mb, an FDR threshold of 0.02, and a clustering radius of 40 kb. The resulting loop calls were used in Fig. 3A. The two genomic loci defining each loop were extracted as loop anchors, which were used in the integration analyses shown in Figs. 3A and 5.

For Fig. S3A, A/B interval counts were divided by compartment genomic coverage and normalized to sum to 100%, excluding mixed and unassigned intervals. For Fig. S3E, proportions were calculated among intervals assigned unambiguously to the eight subcompartments; Expected represents genomic-coverage proportions. For Fig. S3F, positive $\log_2$(observed/expected) values were normalized to sum to one across A1–A4; Expected represents genomic-coverage proportions normalized across A1–A4.

### Public datasets

Processed ini-seq2 (10), DNAscent (11), SNS-seq (12), and BG4 CUT&Tag (13) datasets were obtained from the corresponding published studies and used for integrative analyses.

### OK-seq

FASTQ files from the OK-seq dataset (SRA accession: SRP065949) (14) were downloaded using SRA-Explorer. FASTQ files were processed to generate clean BAM files using the basic processing workflow. OKseqHMM (15) was then used to identify initiation zones (IZs), termination zones (TZs), high flat zones, and low flat zones with the following parameters: thresh=10, chrsizes=hg38.CHROM_SIZES, binSize=1000, and winS=15. The resulting BED files for IZs, TZs, high flat zones, and low flat zones were used for downstream integrative analyses.

### ATAC-seq

FASTQ files (SRR27158649) (16) were downloaded from SRA-Explorer. FASTQ files were processed to generate a narrowPeak file using the basic processing workflow with one modification, i.e. we used "-X 2000" in bowtie2 mapping. The narrowPeak file was used in the integration analysis in Fig. 5.

### Learning 25 chromatin states using ChromHMM

A ChromHMM model was trained to segment the human genome using CUT&RUN and CUT&Tag profiles of ten histone marks, including H2A.Z1, H3K27ac, H3K27me3, H3K36me3, H3K4me3, H3K56ac, H3K9ac, H3K9me3, H4K20me1, and H4K20me3. For each histone mark, multiple biological replicates (≥3, except two for H3K56ac) were available; however, one representative replicate per mark was selected for model training on the basis of the highest FRiP score, as described in the Results. ChromHMM models were learned using multiple state numbers (10, 15, 20, 25, and 30), and the 25-state model with 200bp as bin size was selected as the final configuration, as it provided clear segregation of major chromatin environments while avoiding excessive state fragmentation. This 25-state chromatin state map was subsequently used as a common genomic coordinate framework for downstream integrative analyses, including integration with ORC1 3DM profiles in Fig. 2E, and meta integration analysis with other datasets in Fig. 5.

### Integration of ORC1 3DM with 25 chromatin states

ORC1 3DM was divided into 30 subsets according to peak reproducibility, from eq_1, representing peaks detected in only one library, to eq_30, representing peaks detected in all 30 libraries. These 30 ORC1 peak subsets were used as the x-axis and the 25 chromatin states as the y-axis to generate a heatmap showing the enrichment of ORC1 peaks of different reproducibility across distinct chromatin states. A genomic coverage column was added to indicate the fraction of the genome occupied by each chromatin state. For visualization, the genomic coverage values were normalized by dividing each value by the maximum genomic coverage across the 25 states. Enrichment values were normalized independently for each ORC1 reproducibility subset by dividing the enrichment value for each chromatin state by the maximum value within that subset, scaling each column from 0 to 1.

Group 1 comprised eq_29–30, group 2 eq_22–28, group 3 eq_8–20, and group 4 eq_1–6. The eq_7 and eq_21 subsets were intermediate subsets. Groups 1–3 formed the supergroup, excluding the intermediate subsets. For group-level analyses, selected 50-bp bins were pooled and merged using BEDTools merge -d 100. The all-sites set included eq_1–30.

**Integration of ORC1 3DM, chromatin states, replication initiation sites, and other chromatin contexts**

To represent ORC1 3DM, four subsets of ORC1 peaks (eq_30, eq_26, eq_15, and eq_3) were selected to represent groups 1–4, respectively, of ORC1 binding sites with different levels of reproducibility. Replication initiation datasets included ini-seq2 (15 min and 3 h), DNAscent (HeLa and RPE1 cells), and SNS-seq (core and stochastic initiation sites). Initiation zones, termination zones, high flat zones, and low flat zones were obtained from OK-seq analysis. Additional chromatin features included A/B compartments, TAD boundaries, and loop anchors derived from Hi-C; CTCF-binding sites identified by CTCF ChIP-exo; open chromatin regions identified by ATAC-seq and DNase-seq; and G-quadruplex profiles generated by BG4 CUT&Tag. Genomic coverage of each chromatin state was included as the leftmost column. To visualize relative chromatin-state preferences among datasets generated by different assays, the replication initiation datasets (ini-seq2, DNAscent, SNS-seq, and OK-seq features) and other chromatin datasets (Hi-C features, CTCF ChIP-exo, ATAC-seq, DNase-seq, and BG4 CUT&Tag) were normalized independently across the 25 chromatin states by dividing each value by the maximum within that dataset, scaling values from 0 to 1. To preserve the directly comparable enrichment scale among ORC1 3DM groups and the absolute genomic occupancy of each chromatin state, ORC1 3DM groups and genomic coverage were displayed using their original values.

**Reagents and resources**

| REAGENT or RESOURCE | SOURCE | IDENTIFIER |
|---|---|---|
| Antibodies | | |
| rabbit anti-ORC1 | Bethyl | Cat# A301-892A |
| mouse anti-ORC1 | In-house | 78 1-188 |
| rabbit anti-ORC2 | In-house | CS204 |
| rabbit anti-ORC3 | In-house | CS1890 |
| rabbit anti-ORC4 | In-house | CS2100 |
| rabbit anti-ORC5 | LifeSpan BioScience | Cat# LS-C323686 |
| rabbit anti-ORC6 | MyBioSource | Cat# MBS8247547 |
| rabbit anti-ORCA | Bethyl | Cat# A301-867A |
| rabbit anti-CDC6 | abcam | Cat# ab109315 |
| mouse anti-BM28 (MCM2) | BD | Cat# 610701 |
| rabbit anti-MCM3 | Bethyl | Cat# A300-192A |
| mouse anti-H4K20me1 | Diagenode | Cat# C15200147-10 |
| rabbit anti-H2A.Z1 | Abcam | Cat# ab150402 |
| rabbit anti-H3K27ac | Cell Signaling Technology | Cat# 8173S |
| rabbit anti-H3K27me3 | invitrogen | Cat# MA5-11198 |
| rabbit anti-H3K36me3 | invitrogen | Cat# PA5-96118 |
| rabbit anti-H3K4me3 | Cell Signaling Technology | Cat# 9751S |
| rabbit anti-H3K56ac | Millipore | Cat# 07-677-IS |
| rabbit anti-H3K9ac | Cell Signaling Technology | Cat# 9649T |
| rabbit anti-H3K9me3 | Abcam | Cat# ab176916 |
| rabbit anti-H4K20me3 | Diagenode | Cat# C15410057-10 |
| rabbit anti-HA | Cell Signaling Technology | Cat# 3724S |

| REAGENT or RESOURCE | SOURCE | IDENTIFIER |
|---|---|---|
| rabbit anti-CTCF | Cell Signaling Technology | Cat# 3418S |
| Rabbit mAb IgG isotype control | Cell Signaling Technology | Cat# 3900S |
| Mouse IgG Isotype control | Thermo Fisher | Cat# 10400C |
| Chemicals, peptides, and recombinant proteins | | |
| dTAGv-1 | Tocris | Cat# 6914 |
| BrdU | sigma | Cat# B5002-1g |
| Ampure XP | Beckman coulter | Cat# A63881 |
| CUT&RUN pAG-MNase and Spike-In DNA | Cell Signaling Technology | Cat# 40366 |
| glycogen | sigma | Cat# 10901393001 |
| spermine | sigma | Cat# S3256 |
| Blasticidin S HCl (10 mg/mL) | ThermoFisher | Cat# A1113902 |
| proteinase K | NEB | Cat# P8107S |
| phenol-chloroform-isoamyl alcohol | sigma | Cat# 77617-500ml |
| Biotin-14-dATP | thermofisher | Cat# 19524-016 |
| MboI | NEB | Cat# R0147M |
| T4 DNA Ligase (1 U/µL) | thermofisher | Cat# 15224090 |
| Dynabeads™ MyOne™ Streptavidin T1 | invitrogen | Cat# 65602 |
| Dynabeads™ MyOne™ Streptavidin C1 | invitrogen | Cat# 65001 |
| Benzonase | Sigma | Cat# E8263 |
| MG132 | Calbiochem | Cat# 474790-10MG |
| PhosSTOP | Roche | Cat# 04906845001 |
| IPTG | Roche | Cat# 11411446001 |
| Ni-NTA resin | QIAGEN | Cat# 30210 |
| Amylose resin | NEB | Cat# E8021 |
| Glutathione agarose | Thermo Fisher Scientific | Cat# 16100 |
| Dynabeads Protein G | Thermo Fisher Scientific | Cat# 10004D |
| Azide-PEG3-biotin | Sigma | Cat# 762024 |
| Oligonucleotides | | |
| Listed in Table S1 | | |
| Plasmids | | |
| pX330-1X2 | Addgene | Cat# 58766 |
| pCRIS-PITChv2-BSD-dTAG-BRD4 | Addgene | Cat# 91792 |
| pX330-1X2-ORC1-5-gRNA | This study | N/A |
| pMAL-c2E | Addgene | Cat# 75291 |
| pGEX-6P-1 | Sigma-Aldrich | Cat# GE28-9546-48 |
| pDNR-Dual-FLEX-CTCF | FLEXgene collection | N/A |
| pGEX-6P-CTCF | This study | N/A |
| pMAL-c2E-ORC1 | Stillman lab | N/A |
| pAG/MNase | addgene | Cat# 123461 |
| Bacterial strains | | |
| 5-alpha competent E. coli | NEB | Cat# C29871 |
| BL21 (DE3) competent E. coli | NEB | Cat# C25271 |

| REAGENT or RESOURCE | SOURCE | IDENTIFIER |
|---|---|---|
| Cell lines | | |
| U2OS | ATCC | Cat# HTB-96 |
| U2OS ORC1-FKBP12$^{F36V}$ | This study | |
| Critical commercial assays | | |
| CUTANA™ ChIC/CUT&RUN KIT | EpiCypher | Cat# 14-1048 |
| NEBNext® Ultra™ II End Repair/dA-Tailing Module | NEB | Cat# E7546L |
| NEBNext® Ultra™ II Q5® Master Mix | NEB | Cat# M0544X |
| Lipofectamine™ 2000 Transfection Reagent | invitrogen | Cat# 11668030 |
| Qubit 1X dsDNA HS assay kit | ThermoFisher | Cat# Q33231 |
| Deposited data: Sequencing data have been deposited in GEO under accession GSE351020 and will be released upon publication. | | |
| CUT&RUN, CUT&TAG, ChIP-exo, Hi-C, repli-seq, EdU-seq | This paper | |
| Ini-seq2 | Guilbaud et al. (10) | GEO: GSE186675 |
| DNAscent | Carrington et al. (11) | GEO: GSE265956 |
| SNS-seq | Akerman et al. (12) | GEO: GSE128477 |
| OK-seq | Petryk et al. (14) | SRP065949 |
| ATAC-seq | Hill et al. (16) | GEO: GSM7958836 |
| DNase-seq | Forgetta et al. (17) | GEO: GSM4221655 |
| BG4 CUT&TAG | Hui et al. (13) | GEO: GSE181373 |
| Software and algorithms | | |
| Fastqc v0.12.1 | https://www.bioinformatics.babraham.ac.uk/projects/fastqc/ | RRID:SCR_014583 |
| Trim_galore v0.6.6 | https://www.bioinformatics.babraham.ac.uk/projects/trim_galore/ | RRID:SCR_011847 |
| Cutadapt v5.0 | http://code.google.com/p/cutadapt/ | RRID:SCR_011841 |
| Bowtie2 v 2.4.4 | Langmead and Salzberg (18) | RRID:SCR_005476 |
| Samtools v 1.14 | Li et al. (19) | RRID:SCR_002105 |
| Bedtools v2.30.0 | Quinlan et al. (20) | RRID:SCR_006646 |
| Deeptools v3.5.2 | Ramirez et al. (21) | RRID:SCR_016366 |
| Picard v2.26.11 | Broad Institute, 2019 | RRID:SCR_006525 |
| Macs2 v2.2.9.1 | Zhang and Liu et al. (22) | RRID:SCR_013291 |
| R v4.4.0 | https://www.r-project.org/ | |
| Python v3.13.5 | https://www.python.org | |
| Numpy v1.26.4 | Python package | RRID:SCR_008633 |
| Matplotlib v3.10.5 | Python package | RRID:SCR_008624 |
| Pandas v2.3.2 | Python package | RRID:SCR_018214 |
| Pairtools v1.1.3 | Open2c | RRID:SCR_023038 |
| Cooler v0.10.4 | Open2c | RRID:SCR_024194 |
| Cooltools v0.7.1 | Open2c | RRID:SCR_026118 |
| h5py v3.14.0 | Python package | RRID:SCR_024812 |
| Conda v25.3.1 | https://www.anaconda.com | RRID:SCR_018317 |
| bwa-mem2 v2.3 | Vasimuddin et al. (23) | RRID:SCR_022192 |
| Integrative Genomics Viewer | Robinson et al. (24) | RRID:SCR_011793 |

| REAGENT or RESOURCE | SOURCE | IDENTIFIER |
|---|---|---|
| Juicer v1.22.01 | Durand et al. (25) | RRID:SCR_017226 |
| HiCRep v0.2.6 | Yang et al. (26) | RRID: not available |

**Table S1. Oligonucleotides**

| name | sequence |
|---|---|
| human ORC1 gRNA top strand | 5’-CACCGGTAGTGTGCCATGGCTTCTG-3’ |
| human ORC1 gRNA bottom strand | 5’-AAACCAGAAGCCATGGCACACTACC-3’ |
| forward primer for human ORC1 5’ recombinant donor PCR | 5’-gggccccccctcgagATTGCAGTCTGGCTTTTCCGG-3’ |
| reverse primer for human ORC1 5’ recombinant donor PCR | 5’-ATCGATACCGTCGACGGGAAACCAATTCGCCCAG-3’ |
| forward primer for BSD-P2A-2×HA-FKBP12$^{F36V}$ cassette PCR | 5’-TCCACAGAAGCCATGATGGCCAAGCCTTTGTCTCAAG-3’ |
| reverse primer for BSD-P2A-2×HA-FKBP12$^{F36V}$ cassette PCR | 5’-TGTGGGGTAGTGTGCGCCAGATCCGCCGCCACC-3’ |
| forward primer for human ORC1 5' genotyping PCR | 5′-ATTGCAGTCTGGCTTTTCCG-3′ |
| reverse primer for human ORC1 5' genotyping PCR | 5′-GGCCTGCCAACCCATGAATA-3′ |
| forward primer for human CTCF cloning PCR | 5’-TTCCAGGGGCCCCTGGGATCCATGGAAGGTGATGCAGTC-3’ |
| reverse primer for human CTCF cloning PCR | 5’-GTGATGATGGTGATGGTCgacCCGGTCCATCATGCTGAG-3’ |

**Table S1 (continued). ChIP-exo oligonucleotides**

| name | sequence | |
|---|---|---|
| ExA2_iNN_CTCF_rep1 | [Phos]CAAGCAGAAGACGGCATACGAGATCGTACTAGGTGACTGGAGTTCAGACGTGTGCTCTTCCGATCT | first adapter top strand |
| ExA2_iNN_CTCF_rep2 | [Phos]CAAGCAGAAGACGGCATACGAGATTAGGCATGGTGACTGGAGTTCAGACGTGTGCTCTTCCGATCT | first adapter top strand |
| ExA2_iNN_CTCF_rep3 | [Phos]CAAGCAGAAGACGGCATACGAGATCGAGGCTGGTGACTGGAGTTCAGACGTGTGCTCTTCCGATCT | first adapter top strand |
| ExA2_iNN_CTCF_rep4 | [Phos]CAAGCAGAAGACGGCATACGAGATATCTCAGGGTGACTGGAGTTCAGACGTGTGCTCTTCCGATCT | first adapter top strand |
| ExA2_iNN_CTCF_rep5 | [Phos]CAAGCAGAAGACGGCATACGAGATCGGAGCCTGTGACTGGAGTTCAGACGTGTGCTCTTCCGATCT | first adapter top strand |
| ExA2_iNN_CTCF_rep6 | [Phos]CAAGCAGAAGACGGCATACGAGATACTGAGCGGTGACTGGAGTTCAGACGTGTGCTCTTCCGATCT | first adapter top strand |
| ExA2_iNN_IgG_rep1 | [Phos]CAAGCAGAAGACGGCATACGAGATTAAGGCGAGTGACTGGAGTTCAGACGTGTGCTCTTCCGATCT | first adapter top strand |
| ExA2_iNN_IgG_rep2 | [Phos]CAAGCAGAAGACGGCATACGAGATGGACTCCTGTGACTGGAGTTCAGACGTGTGCTCTTCCGATCT | first adapter top strand |
| ExA2_iNN_IgG_rep3 | [Phos]CAAGCAGAAGACGGCATACGAGATGCTACGCTGTGACTGGAGTTCAGACGTGTGCTCTTCCGATCT | first adapter top strand |
| ExA2_iNN_IgG_rep4 | [Phos]CAAGCAGAAGACGGCATACGAGATGCTCATGAGTGACTGGAGTTCAGACGTGTGCTCTTCCGATCT | first adapter top strand |
| ExA2_iNN_IgG_rep5 | [Phos]CAAGCAGAAGACGGCATACGAGATGCGTAGTAGTGACTGGAGTTCAGACGTGTGCTCTTCCGATCT | first adapter top strand |
| ExA2_iNN_IgG_rep6 | [Phos]CAAGCAGAAGACGGCATACGAGATTAGCGCTCGTGACTGGAGTTCAGACGTGTGCTCTTCCGATCT | first adapter top strand |
| ExA2B | GATCGGAAGAGCACACGTCTGAACTCCAGTCAC | first adapter bottom strand |
| ExA1-58 | AATGATACGGCGACCACCGAGATCTACACTCTTTCCCTACACGACGCTCTTCCGATCT | second adapter top strand |
| ExA1-SSL_N5 | NNNNNAGATCGGAAGAGCG | second adapter bottom strand |
